# Global multi-environment resistance QTL for foliar late blight resistance in tetraploid potato with tropical adaptation

**DOI:** 10.1101/2020.02.16.950618

**Authors:** Hannele Lindqvist-Kreuze, Bert de Boeck, Paula Unger, Dorcus Gemenet, Xianping Li, Zhechao Pan, Qinjun Sui, Junhong Qin, Gebremedhin Woldegjorgis, Kassaye Negash, Ibrahim Seid, Betaw Hirut, Manuel Gastelo, Jose De Vega, Merideth Bonierbale

## Abstract

The identification of environmentally stable and globally predictable resistance to potato late blight is challenged by the crop’s clonal and polyploid nature and the pathogen’s rapid evolution. Genome-wide analysis (GWA) of multi-environment trials can add precision to breeding for complex traits. A diversity panel of tetraploid potato germplasm bread for multiple resistance and quality traits was genotyped by genotyping by sequencing (GBS) and phenotyped for late blight resistance in a trait observation network spanning three continents addressed by the International Potato Center’s (CIP’s) breeding program. The aims of this study were to (i) identify QTL underlying resistance in and across environments and (ii) develop prediction models to support the global deployment and use of promising resistance sources in local breeding and variety development programs. Health-indexed *in vitro* plants of 380 clones and varieties were distributed from CIP headquarters in Peru to China and Ethiopia and tuber seed was produced centrally in each country. Phenotypes were recorded as rAUDPC following field exposure to local isolates of *Phytophthora infestans*, Stringent filtering for individual read depth >60 resulted in 3,239 tetraploid SNPs. Meanwhile, 55,748 diploid SNPs were identified using diploidized data and individual read depth>17. The kinship matrix was utilized to obtain BLUP and identify best performing germplasm in each and all environments. Genotypes with high levels of resistance in all environments were identified from the B3, LBHT and B3-LTVR populations. GWA identified stable QTL for late blight resistance in chromosome 9 and environment specific QTL in chromosomes 3, 5, 6 and 10.

## Introduction

Potato genetic resources comprise a polyploid series consisting of a tremendously diverse germplasm of wild relatives and cultivated landraces (Spooner 2014; Ovchinnikova et al., 2011). However, most commercially cultivated potato varieties are tetraploid (2n=4x=48) with the genome consisting mostly of *Solanum tuberosum* Group *tuberosum* with some introgressions from a few wild species and cultivated landraces (Bradshaw et al., 2006; reviewed by Bethke et al., 2017; reviewed by Gaiero et al., 2018). Tetraploid potato is a highly heterozygous, outcrossing autopolyploid, crop which complicates genetic analysis. Most of the early genetic mapping studies utilized bi-parental populations at the simpler, diploid level (2n=2x=24) and several disease resistance loci were identified in the genome of potato this way (reviewed by Gebhardt and Valkonen 2001). However, this approach does not permit the assessment of large gene pools or multi-allelic interactions that influence traits in polyploids. Significant progress has recently been made in the development of algorithms and software for genotype calling, linkage and QTL analysis in polyploid species. SNP arrays have been developed for potato: 8K SolCAP (Hamilton et al., 2011) and the 20K SolSTW arrays (Vos et al., 2015). These were developed using North American and European potato germplasm, respectively, and are not consequently the best options for genotyping CIP germplasm since it contains more introgressions from the native South American gene pool. According to our previous experience, less than 50% of the SNPs on the 8K SolCAP array were informative in a test sample of CIP germplasm (Lindqvist-Kreuze et al., 2014). Genotyping by sequencing (GBS) has been applied to tetraploid potato (Uitdewilligen et al., 2013, Sverrisdottir et al., 2017); and variant calling from short read sequencing data considering allele dosage is now possible using several different tools, such as GATK, Freebayes, or SAMtools to name a few (Clevenger et al., 2015). However, reliable dosage calling in the heterozygous individuals depends on the read depth in the SNP loci. It was recently demonstrated in autopolyploid blueberry, that a read depth of 61 was adequate to reliably call the allele dosage, while only 17 reads were needed to reliably classify simplex tetraploids as heterozygous with 95% accuracy (Matias et al., 2019). The identification of QTL in autopolyploids is facilitated by new tools, such as called GWASpoly that considers allele dosage effects (Rosyara et al., 2016). Together, these advances make genomic analysis of tetraploid potato more informative and applicable to evolutionary and breeding studies.

The goal of CIP potato breeding program is to develop resilient, high yielding, healthy and early maturing varieties for small-holder farming systems in the developing world. We are targeting farming systems that usually function with minimum input of pesticides and therefore a high level of disease resistance is an indispensable trait. To this end, CIP’s potato breeding program has developed breeding populations selected for high levels of resistance to late blight caused by the oomycete *Phytophtora infestans*, and resistance to Potato Virus Y (PVY), Potato Virus X (PVX) and Potato Leaf Roll Virus (PLRV). Previous studies have identified genomic regions in CIP’s breeding germplasm explaining resistance to late blight focusing on phenotypic data collected from field trials in Peru or using local pathogen strains in greenhouse conditions (Li et al., 2010; Lindqvist-Kreuze et al., 2014, Jiang Rui et al., 2018). Performance information has been sporadically published about CIP’s bred materials in the target regions where they have been distributed to (Muhinyuza et al., 2015; Hirut et al., 2017b) but to our knowledge no genetic analysis has been published identifying resistance QTL for resistance in CIP germplasm tested in environments outside Peru.

The overall goal of this research was to collect data on the foliar late blight resistance of CIP’s advanced tetraploid potato clones in Ethiopia and China to inform breeding decisions. To systematically evaluate CIP’s breeding materials in diverse target environments we established a trait observation network (TON) of collaborators and assembled a diversity panel that consists of representative advanced clones (including elite materials) from each of CIP’s breeding populations. This so-called TON panel was then distributed from Peru to China and Ethiopia, where it was included in a series of trait evaluation experiments by national research and CIP institutions. The specific aims aims of this study were to (i) identify QTL underlying resistance in and across environments and (ii) develop prediction models to support the global deployment and use of promising resistance sources in local breeding and variety development programs.

We report the genotyping, estimation of linkage disequilibrium and population structure of the TON panel and identification of QTL for late blight resistance via genome wide association (GWA). In addition, we present a case for genomics assisted breeding for foliar late blight resistance and show how the use of genomics and pedigree information can be used to select best bet clones for breeding and variety development in diverse target environments.

## Materials and methods

### Germplasm

The TON panel used in this study consisted of 380 genotypes representing seven CIP breeding populations as well as a group of varieties with variable origins (Table 1.). ‘Population A’ was developed at 1980-1990 with emphasis on late blight resistance. Sources of late blight resistance were improved materials with *S. demissum*-derived resistance from breeding programs around the world, native Andean cultivars *S. tuberosum* groups *andigena*, *phureja* and *stenotomun*, wild species *S. acaule* and *S. bulbocastanum*. ‘Population B3’ genotypes were derived from ‘Population A’ with emphasis on increasing frequencies and levels of quantitative resistance to late blight. The ‘B1 population’ is derived from *S. tuberosum* group *andigena*. The ‘LTVR population’ is characterized mainly for its resistance to the most important virus diseases (PVY, PVX and PLRV), short crop duration, and adaptation to warm environments. The ‘LB-HT’ population combines late blight resistance from the ‘B3 population’ and heat tolerance from North American and European bred varieties and the LTVR population. ‘B3-LTVR’ population contains hybrid genotypes originating from crosses between ‘B3’ and ‘LTVR populations’. The ‘pre-Bred’ population has genotypes that have LB resistance introduced from wild *Solanum* species into the tetraploid background of ‘B3’ or ‘LTVR’. The varieties group consists of a group of potato varieties or key breeding lines: ‘Desiree’, ‘Atlantic’, ‘Spunta’, ‘Granola’, ‘Yungay’, ‘Tomasa Condemayta’, ‘DTO-33’, ‘Kufri Yoti’, and ‘Chucmarina’. CIP numbers and the parentage of the 380 genotypes are given in the Supplementary Table S1.

**Table 1.** Contribution of CIP breeding populations to the TON diversity panel

| **Breeding population** | **Genotypes evaluated** | **main breeding objective** |
| --- | --- | --- |
| A | 13 | late blight resistance |
| B1 | 11 | late blight resistance |
| B3 | 100 | late blight resistance |
| B3-HT | 37 | late blight resistance, heat tolerance |
| B3-LTVR | 25 | Hybrid population combining late blight resistance, heat tolerance, virus resistance |
| LTVR | 186 | virus resistance, heat tolerance, drought tolerance, salinity tolerance |
| PREBRED | 2 | late blight resistance |
| VARIETY | 6 | varied |
| Grand Total | 380 |  |

### Environments

The field sites in Ethiopia and China are important potato production areas, while in the field site in Peru, potato is not the main crop (Table 2). The late blight pathogen populations have been described in each location. In Peru and Ethiopia only the A1 mating type has been identified and different clonal lineages are present that frequently contain virulence to most of the known *S. demissum R* genes (Lindqvist-Kreuze et al.; 2019 Mihretu et al., 2019). In contrast in Southern China A2 mating type has been found dominating (Chen et al., 2017).

**Table 2.**
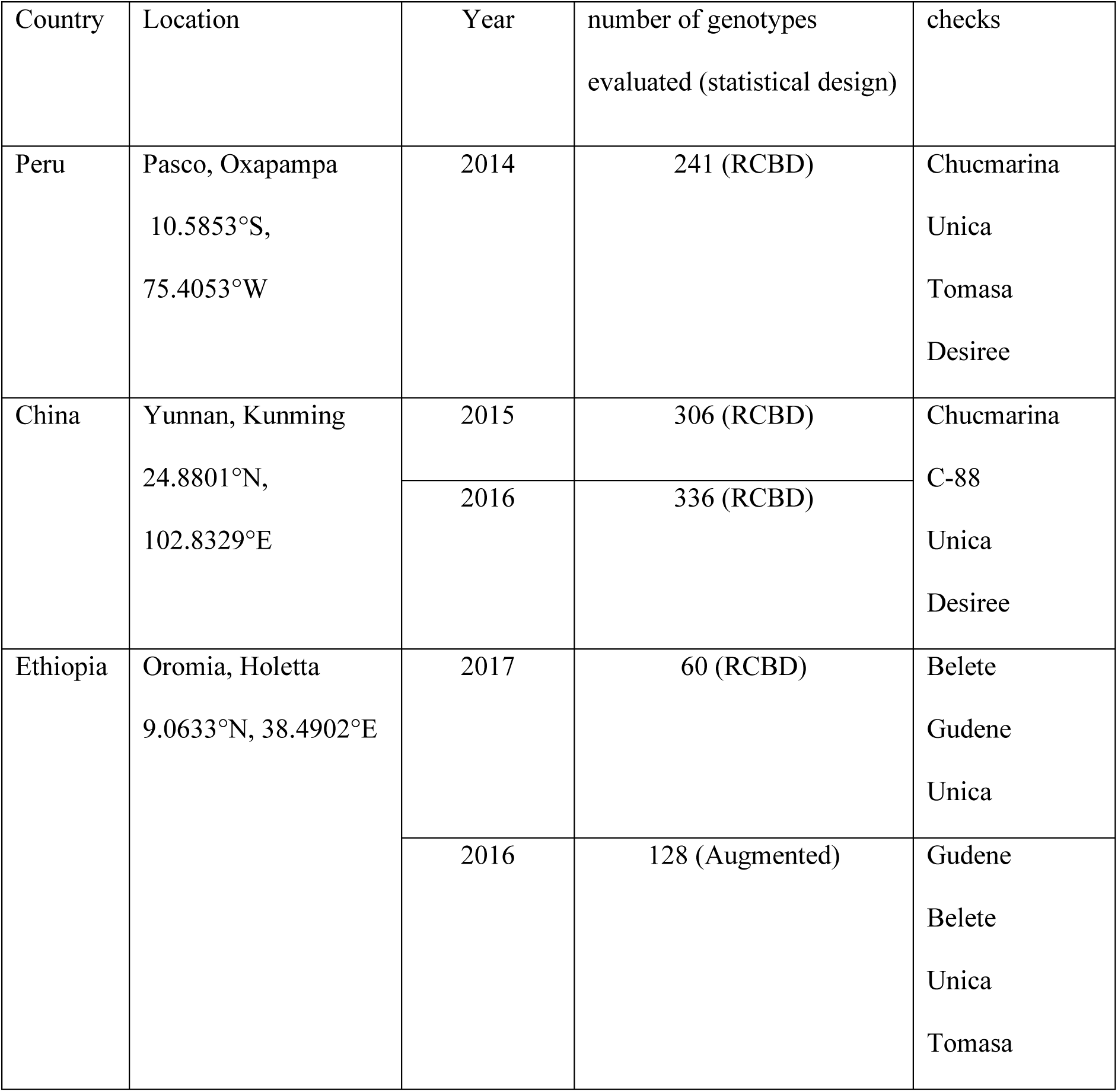
Description of the phenotypic evaluations involving the TON panel clones.

### Field trials

Standard protocols at CIP were utilized for planning and conducting the field trials (Bonierbale, 2007). The statistical designs in each trial are shown Table 2. Uniform tuber seed was produced centrally in each country following the introduction of *in vitro* plants or mini-tubers from CIP facilities in Peru or Kenya.

Late blight resistance was evaluated under endemic disease pressure. The disease level in the plots was recorded as ‘percent leaf area infected’ at 7-day intervals until susceptible controls reached 100% infection. These values were used to calculate the area under the disease progress curve (AUDPC) and relative AUDPC (rAUDPC). The data was collected and processed using the HiDAP field book system (https://research.cip.cgiar.org/gtdms/hidap/).

### Statistical analysis of phenotypic data

From the weekly observations of the disease incidence in the plots, the AUDPC was calculated and the estimated means (BLUEs) were transformed to the relative AUDPC (rAUDPC) to facilitate the comparisons among the different locations. The best linear unbiased predictor (BLUP) and best linear unbiased estimator (BLUE) and values as well as ranked predictors were calculated using ASREML package.

### Genotyping, variant calling and filtering for association analysis

In total 380 potato clones were genotyped. Library construction and genotyping by genotyping by sequencing (GBS) was outsourced to the Genomics Facility at Cornell University in 2015. The DNA was digested with EcoT221 restriction enzyme and the libraries were 48x multiplexed for sequencing. The diploid calling was by the service provider using the Tassel pipeline (Bradbury et al., 2007). The resulting Variant Call Format (VCF) file was processed with Bcftools (https://samtools.github.io/bcftools/) to filter the variants for minimum read depth (RD) of 17, minimum genotype quality (GQ) of 30, and minor allele frequency (MAF) 0.03. The SNPs that didn’t pass these criteria were changed to missing call, and finally only the SNP sites that contained less than 30% missing data were selected.

For polyploid calling the raw FASTQ files were processed with Stacks (Catchen et al., 2013) to remove the barcodes and with TrimGalore https://github.com/FelixKrueger/TrimGalore to trim the ends of reads. The reads were aligned to the reference genome version *S. tuberosum_448_v4.03* (Sharma et al., 2013) using BWA (Li and Durbin, 2009) and the resulting SAM files were converted to BAM files using Samtools (Li et al., 2009). The variants were called using GATK HaplotypeCaller option (Poplin et al., 2017), disabling the duplicate read filter (this is recommended for GBS data) and joint genotyping using the -ERC GVCF mode. From the VCF files SNP calls were filtered using Bcftools for minimum RD of 61, minimum GQ of 30 and MAF of 0.03. The samples that didn’t pass these criteria were changed to missing call, and finally only the SNP sites that contained less than 30% missing data were included in the analysis.

### Analysis of the Population sub-structuring

Population sub-structuring with tetraploid data was done using PolyRAD (Clark et al., 2019). Only variants in pairwise LD under 0.1 were previously filtered (LD pruning). The diploid dataset was analyzed using SNPrelate (Zheng et al., 2012) using the subset of bi-allelic SNPs, filtering for LD (0.2) MAF (<0.03) and missingness (<0.3).

### GWA

Marker trait associations were modelled for all the trials independently and using the diploid and the tetraploid marker sets with the GWASpoly package (Rosyara et al., 2016). For the tetraploid dataset *general, additive, simplex dominant* (1-dom) and *duplex dominant* (2-dom) models were used while for the diploid dataset *diplo-general, diplo-additive*, and the *simplex dominant* (1-dom) model were used. The parameters used for the GWAS modelling function *GWASpoly* in R were the following: no fixed effects, 4 principal components were included as covariates, a minimum MAF of 0.03 and a maximum genotype frequency (after applying dominance relations) equal to 0.95 were set, and P3D approximation was used. To detect statistical significance, the Bonferroni correction method was used, ensuring the genome-wide type I error is not greater than 0.05. Manhattan plots were generated to display significant SNP in the different genetic models. In addition, Q-Q plots were used to evaluate the goodness of fit of the genetic model and the quality of the phenotypic data.

The genomic positions of the resulting SNPs associated with plausible QTL for pathogen resistance in relation to other loci, known genes and QTL, were determined using the *S. tuberosum* Group Phureja DM1-3 516R44 (v4.03) pseudomolecule browser (http://solanaceae.plantbiology.msu.edu/) available from the Potato Genome Sequence Consortium (PGSC). To obtain an approximate of the physical location of markers for pathogen resistance present in literature, the position in cM was obtained from the GABI Primary Database (https://www.gabipd.org/projects/Pomamo/) and then “translated” to an approximate physical position in Mbp using information provided in Sharma et al. (2013) that integrates the potato genome and physical and genetic maps.

## Results and discussion

### Population structure

The tetraploid dataset included a total of 305,345 SNPs after the GATK variant calling, while the diploidized dataset after Tassel pipeline had a SNP count of 312,727. After filtering these datasets, the Principal component analyses based on 23,804 diploid SNPs and 182,435 tetraploid variants identified no strong population sub-structuring (Figure 1 for the tetraploid data, Figure S1 for the diploid data) in the diversity panel which makes it an ideal genotype set for GWA. Only population ‘B1’ separates from the rest most likely because the genetic background of the ‘B1’ population is *S. tuberosum* group *andigena*, while the rest are mostly group tuberosum type. ‘LB-HT’ shares alleles with the ‘B3’ population, as expected since these clones are hybrids with B3 clones in their pedigrees. ‘B3’ and ‘LTVR’ population clones are also mostly separated with a few exceptions of clones that may have been mislabelled. ‘B3-LTVR’, which is a hybrid between the two populations and this can be clearly seen in the PCA plot as well. Not surprisingly, Population ‘A’ is intermingled within population ‘B3’ since the ancestors of the ‘B3’ clones were selected clones of the ‘A’ population.

**Figure 1.**
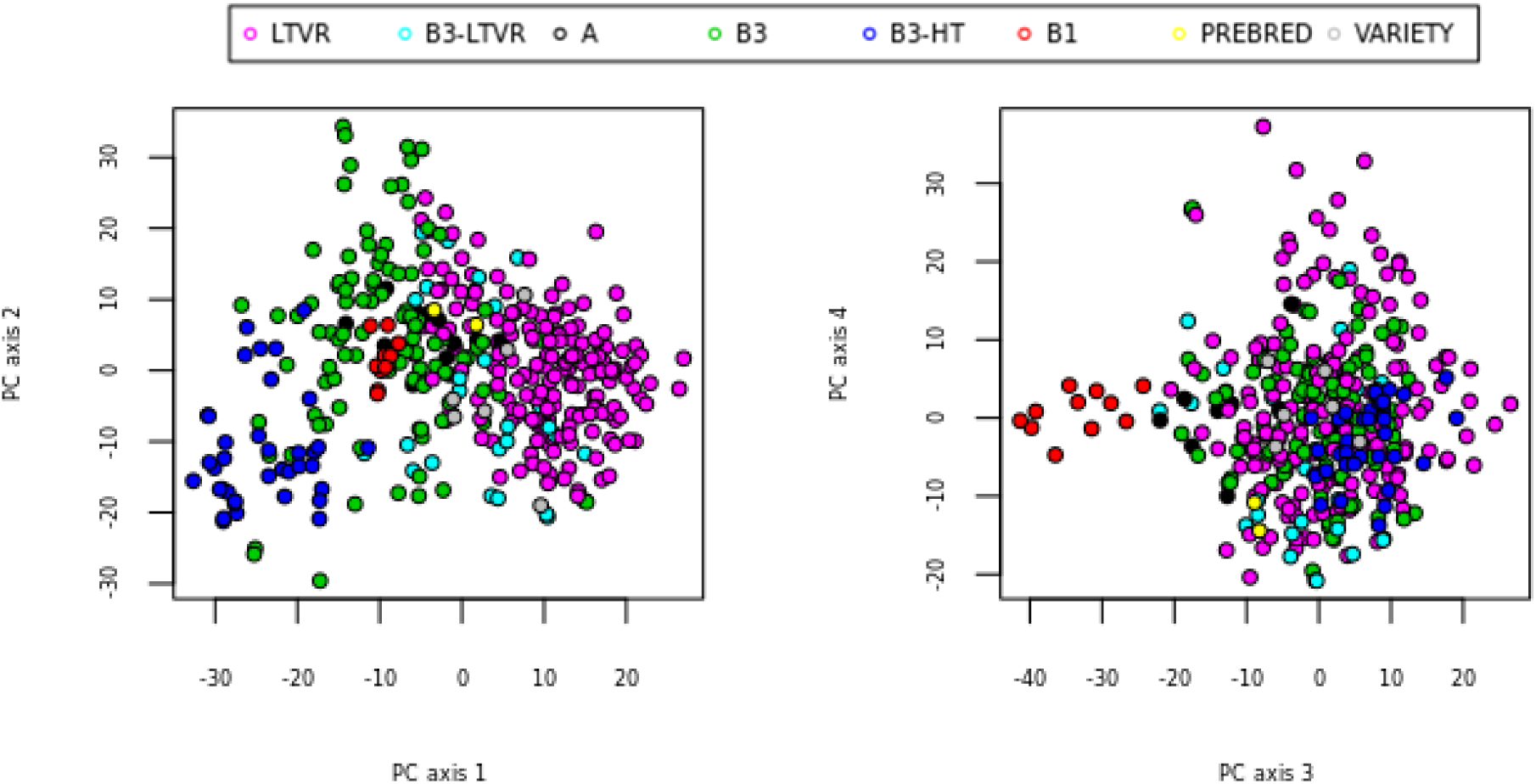
Population sub-structuring based on polyRAD estimation of genotype probabilities

### SNP sets for GWA

After applying filtering parameters tailored for GWA to all variants in both datasets, the numbers of SNPs were reduced to 3,239 tetraploid SNPs and 55,748 diploid SNPs. The DP thresholds were based on the Matias et al. 2019. The study points out, that assuming the GBS method entails 0.5% allelic error, a minimal RD of 17 is necessary in order to classify simplex tetraploid calls as heterozygous with a 95% accuracy. To annotate the allele dosage with the same accuracy, a much higher RD of 61 is needed. Both sets of filtered SNPs were used in the GWAS to identify trait-linked QTL.

### Linkage disequilibrium

LD decay was estimated using the tetraploid marker set. A spline was fitted on the 90^th^ percentile of the squared correlation coefficient between the alleles in a pair of markers (r^2^) and the physical distance between these pairs of markers on the “short” distance of up to 10 Mb (Figure 2A) and “long” distance up to 80 Mb (Figure 2B) over all chromosomes. The intersection of a defined significance threshold of r^2^=0.1 and the fitted spline allowed us to estimate the short distance vs long distance LD decay. On the short distance the threshold is reached at 2Mb, while on the long distance it is reached at 5.5Mb. Considering the short distance LD-decay estimate of r^2^ _1/2max, 90_, which was suggested as the most consistent estimator for LD decay in potato by Vos et al (2017), we obtain the r^2^ _1/2max, 90_ value of 0.55 Mb. This r^2^ _1/2max, 90_ value is equivalent in Vos (2017) data for recent European potato varieties (0.6 Mb) and a bit lower than the study of Sharma et al., (2018) where the r^2^ _1/2max, 90_ value was 0.91 Mb. The average r^2^ for the short distance in our dataset was 0.091, which is a bit lower than the average r^2^ (0.19-0.22) reported for the European varieties (Vos et al., 2017), indicating that there were probably more founder haplotypes in our diversity panel than in the European pooled varieties. The LD decay estimated was moderate, and comparable to the LD decay found in the European potato germplasm. Estimates based on the average r^2^ of the markers along the short distance suggest that high diversity is retained in the germplasm and that tens of thousands of markers would be needed to cover the entire tetraploid genome.

**Figure 2.**
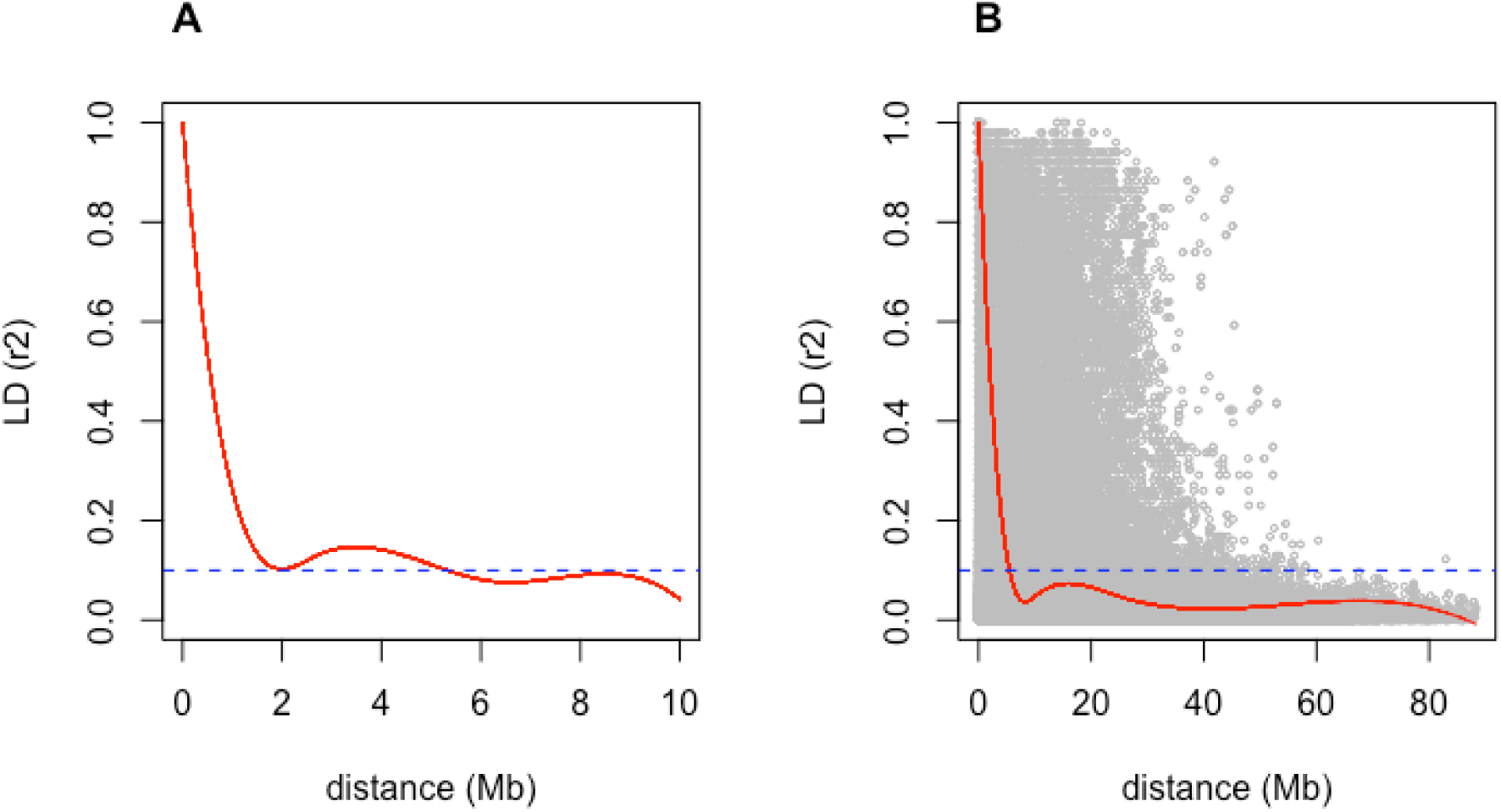
Linkage disequilibrium (LD) estimated in the TON panel based on Pearson correlation coefficient (r^2^) plotted against the physical map distance (Mb) between pairs of SNP.

### Late blight resistance

There was a high number of genotypes with rAUDPC values comparable to the resistant control genotype which is released as a variety called Chucmarina in Peru and as Belete in Ethiopia (Figure 3). Notably, most of the genotypes tested in China were more resistant than the local variety C-88 that has been popular because of its good late blight resistance.

**Figure 3.**
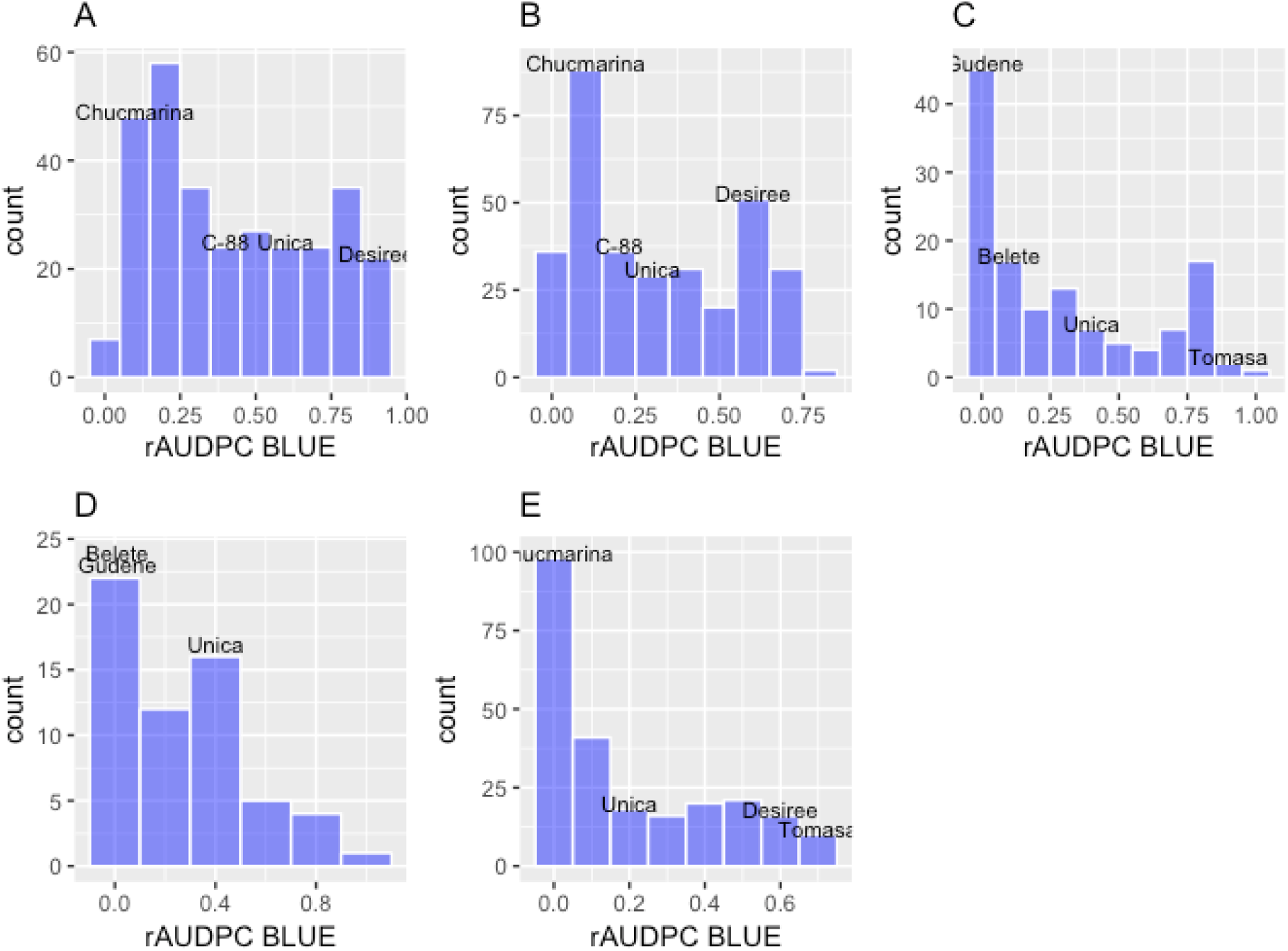
Histograms of rAUDPC values in Kunming, China in 2015(A) and in 2016 (B), Holeta, Ethiopia in 2016 (C) and 2017 (D), and Oxapampa, Peru at 2014 (E). The control genotypes (checks) in each trial are indicated in the plots based on their rAUDPC value.

In this research project over 300 advanced tetraploid clones from CIP were shared with partners, but due to various reasons not all were evaluated in all environments. The genotypes can only be internationally distributed as *in vitro* plants with a health certificate. After receiving the plants, there needs to be at least two rounds of multiplication involving either cuttings or tubers to obtain seed for the replicated trials. However, the performance of all genotyped clones and their pedigree parents could be estimated in all environments by incorporating the marker data into the mixed model.

The GGE biplot for predicted values (BLUP) and marker-based kinship matrix shows the performance of the genotypes and some of their parents in all environments (Figure 4). The most resistant genotypes belong to the B3 and B3-HT populations, while only a few from the LTVR and B3-LTVR had high level of resistance. From this figure, we evidenced that some genotypes’ resistance to late blight is environment specific, nevertheless several genotypes show stable resistance across environments. The correlations among environments were high (Figure 5). Particularly the environment in Peru is highly correlated with all the other environments suggesting that resistant clones selected in Peru will also likely have good resistance in these other environments.

**Figure 4.**
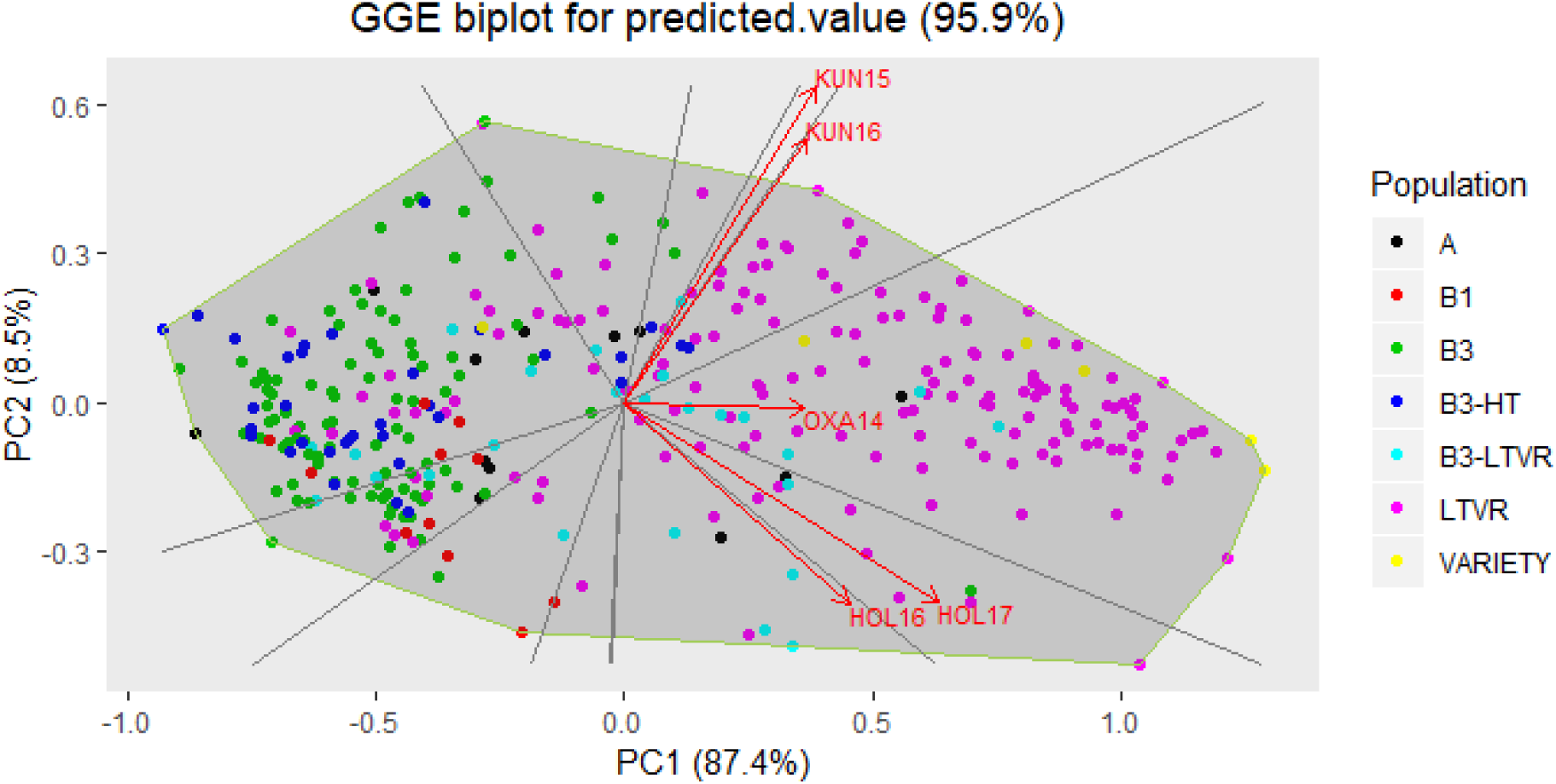
GGE biplot for predicted performance (based on BLUP and genetic kinship matrix) of the test genotypes and their parents in Kunming 2015 (KUN15) and 2016 (KUN16), Oxapampa 2014 (OXA14), Holetta 2016 (HOL16) and 2017 (HOL17).

**Figure 5.**
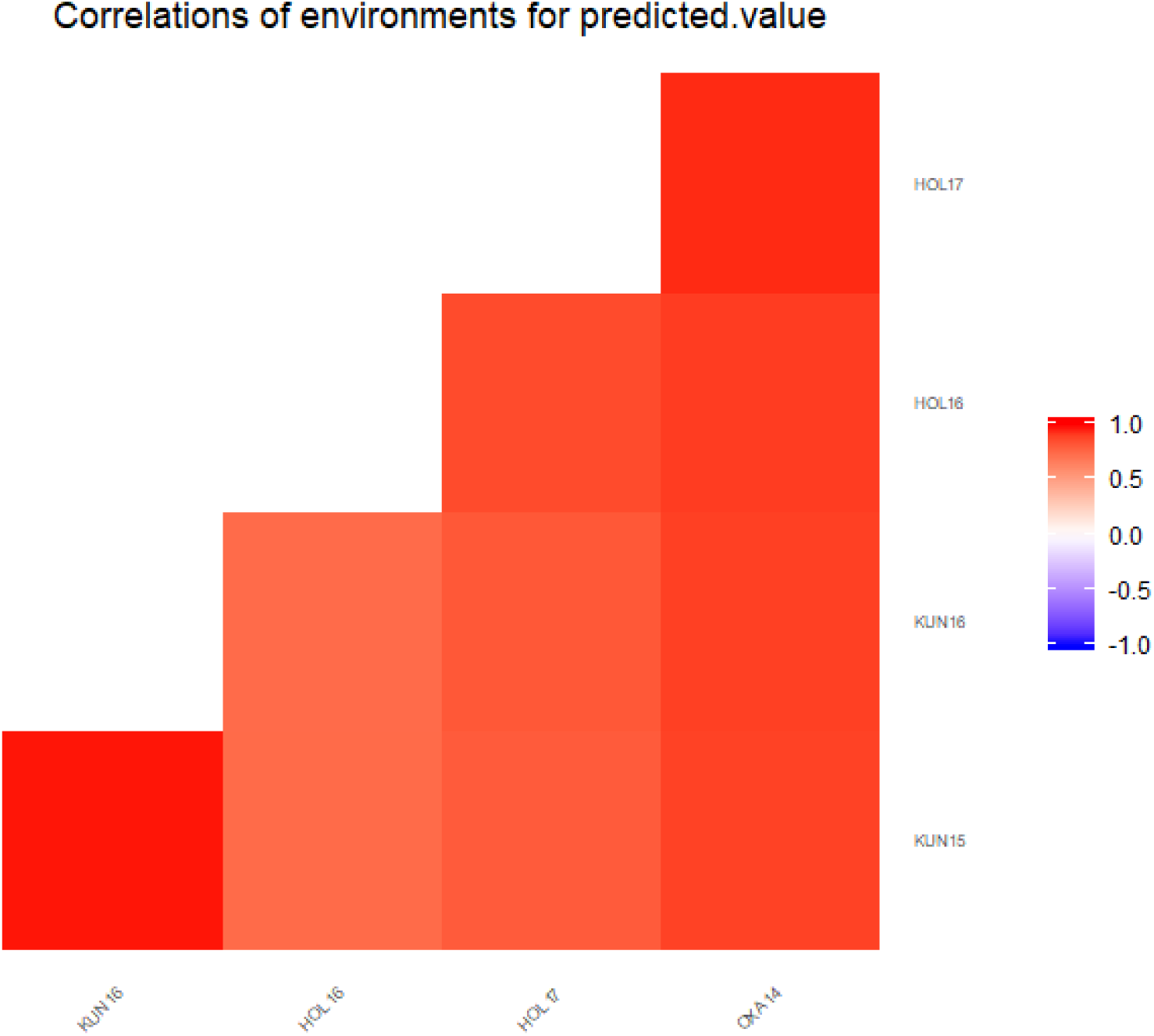
Heat-plot on the correlations among the different trials.

CIP’s breeding strategy defined in the 1990s focused on improving the quantitative resistance in the B3 population by phenotypic recurrent selection under endemic pressure from the “new population” of *P. infestans* in the Peruvian Andes supplemented by progeny tests to identify parents with good general combining. ability and eliminate those resulting in segregation for hypersensitive response against the test isolates. The pathogen population in this area is dominated by the A1 mating type and EC-1 clonal lineage, which is highly aggressive and complex in its virulence (Lindqvist-Kreuze et al., 2019). Despite the differences in the pathogen populations in terms of the mating type and clonal lineages among the countries it seems that phenotypic selection for late blight resistance in Peru was largely successful and results transferable across the three environments tested here.

### QTL for late blight resistance

Several SNPs were significantly associated with late blight resistance in the field trials with a total of 16 markers tagging possible QTL (Table 3 and Table 4). In the tetraploid data, 6 markers for late blight resistance were found in chromosomes III, V and IX while in the diploid dataset 14 markers on chromosomes 0, III, V, VI, IX and X were associated with the resistance phenotype. Populations of *P. infestans* are diverse and there is a trend of increasing diversity in potato-growing regions worldwide (Cooke and Lees, 2004). Taking this into account, the markers on chromosome IX could indicate a QTL for broad resistance not specific to regional late blight strains, because the QTL was observed in data from trials in Peru and China, while markers associated to the QTL on chromosomes III, V and VI were found with the data of unique locations (Table 3 and Table 4). For example, the neighbouring SNPs on chromosome III, could be indicating a QTL particularly responsible for resistance against late blight strains specific to Holeta, Ethiopia. On the other hand, the marker on chromosome X found in the diploidized data could indicate a QTL for broad resistance having been observed in data collected in Peru as well as in China (Table 4). The highest number of SNPs associated to late blight resistance in the GWAS were mapped between 59 and 61.2 Mbp of chromosome IX. All mapped to the end of the long arm of the chromosome, in a region that had been previously associated with late blight resistance in Peru (Lindqvist-Kreuze et al., 2014). Additionally, the markers were within or surrounding the segment between 59.3 and 61.0 Mbp that forms a large cluster of putative resistance genes. For example, the locus PGSC0003DMG400020587 encodes a homolog of *Rpi-vnt1* (Mosquera et al., 2015), a major gene for resistance to *P. infestans* that has been previously cloned and characterized in the wild potato species *Solanum venturii* (Pel et al., 2009; Foster et al., 2009).

**Table 3.** Markers tagging QTL for late blight resistance with the general, additive and 1-dominance models of GWASpoly and tetraploid SNP.

| Marker (chromosome followed by position) | Ref | Alt | Trait | Model | Threshold | Score | Effect |
| --- | --- | --- | --- | --- | --- | --- | --- |
| 0_36073482 | G | A | Oxa2014 | general | 4.71 | 8.64 | NA |
|  |  |  |  | additive | 4.78 | 8.64 | -0.18 |
|  |  |  |  | 1-dom-alt | 4.71 | 8.64 | -0.18 |
|  |  |  | Yun2015 | general | 4.74 | 20.21 | NA |
|  |  |  |  | additive | 4.78 | 20.21 | -0.29 |
|  |  |  |  | 1-dom-alt | 4.71 | 20.21 | -0.29 |
|  |  |  | Yun2016 | general | 4.75 | 19.37 | NA |
|  |  |  |  | additive | 4.78 | 19.37 | -0.25 |
|  |  |  |  | 1-dom-alt | 4.71 | 19.37 | -0.25 |
| 3_3319097 | A | T | Hol2017 | general | 4.49 | 5.83 | NA |
|  |  |  |  | 1-dom-alt | 4.71 | 6.05 | 0.43 |
| 5_4260524 | A | T | Yun2016 | 1-dom-alt | 4.71 | 5.01 | 0.14 |
| 9_58779951 | G | A | Oxa2014 | additive | 4.78 | 4.84 | -0.11 |
|  |  |  |  | 1-dom-alt | 4.71 | 5.52 | -0.13 |
|  |  |  | Yun2015 | general | 4.74 | 4.94 | NA |
|  |  |  |  | 1-dom-alt | 4.71 | 5.92 | -0.14 |
|  |  |  | Yun2016 | 1-dom-alt | 4.71 | 4.77 | -0.11 |
| 9_59967523 | A | T | Yun2015 | general | 4.74 | 7.3 | NA |
|  |  |  |  | additive | 4.78 | 6.94 | -0.14 |
|  |  |  |  | 1-dom-alt | 4.71 | 8.18 | -0.17 |
|  |  |  | Yun2016 | general | 4.75 | 7.76 | NA |
|  |  |  |  | additive | 4.78 | 8.29 | -0.13 |
|  |  |  |  | 1-dom-alt | 4.71 | 8.38 | -0.15 |
| 9_60067335 | A | G | Hol2016 | 1-dom-alt | 4.71 | 4.81 | -0.22 |
|  |  |  | Oxa2014 | general | 4.71 | 11.11 | NA |
|  |  |  |  | additive | 4.78 | 11.64 | -0.2 |
|  |  |  |  | 1-dom-alt | 4.71 | 12.1 | -0.21 |
|  |  |  | Yun2015 | general | 4.74 | 19.39 | NA |
|  |  |  |  | additive | 4.78 | 18.99 | -0.26 |
|  |  |  |  | 1-dom-alt | 4.71 | 20.31 | -0.28 |
|  |  |  | Yun2016 | general | 4.75 | 16.53 | NA |
|  |  |  |  | additive | 4.78 | 17.26 | -0.22 |
|  |  |  |  | 1-dom-alt | 4.71 | 17.59 | -0.23 |

**Table 4.**
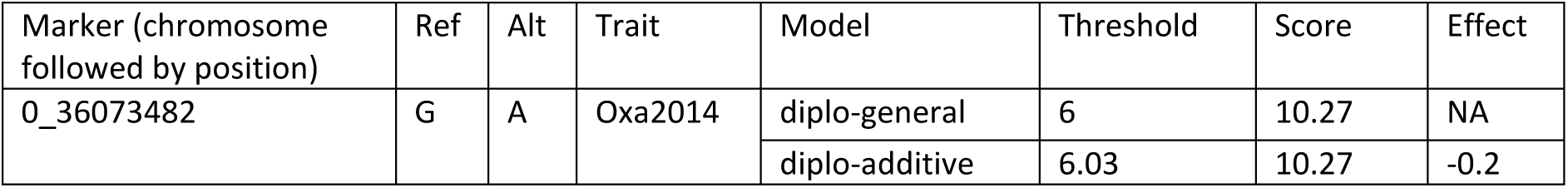
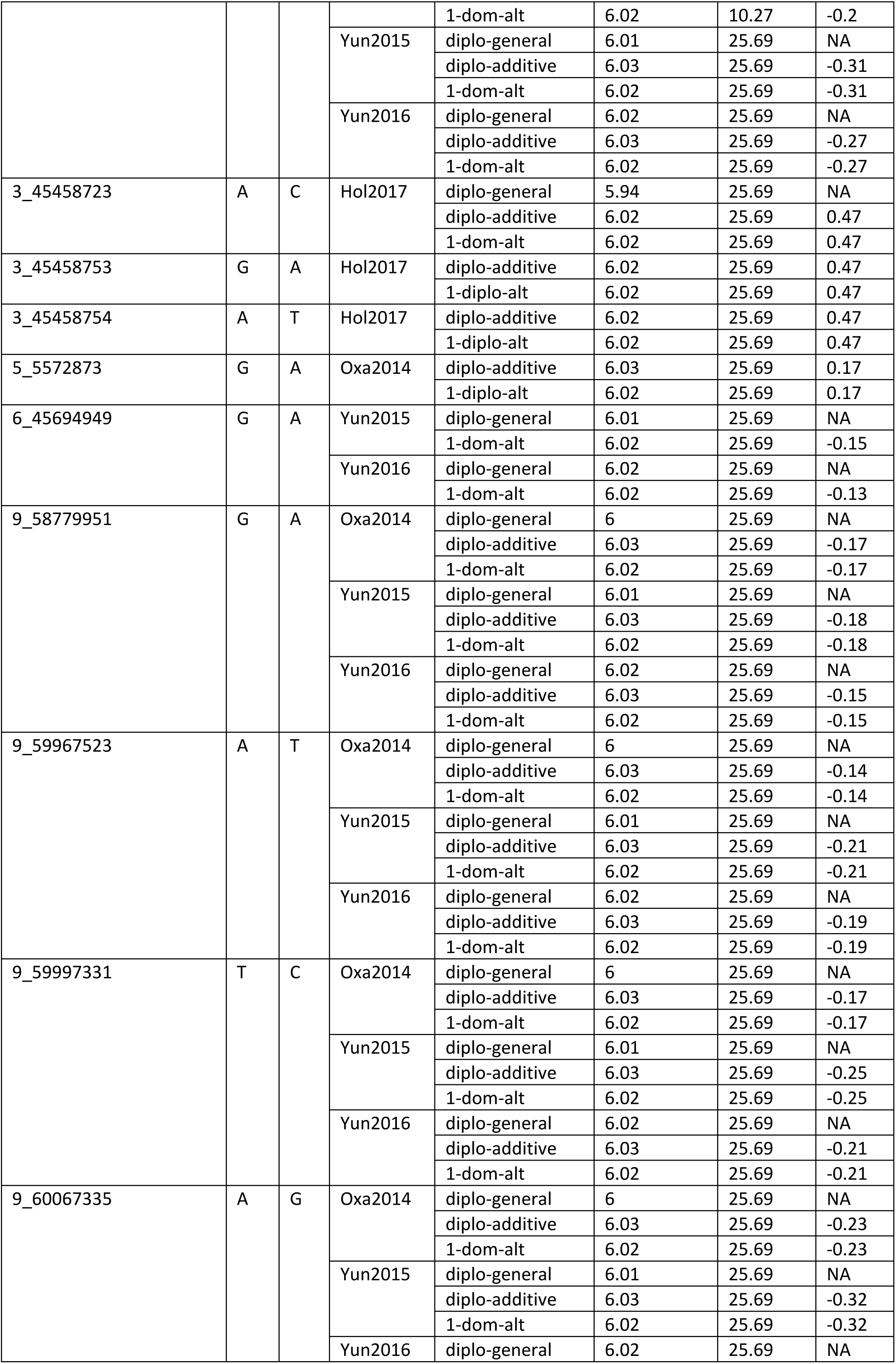
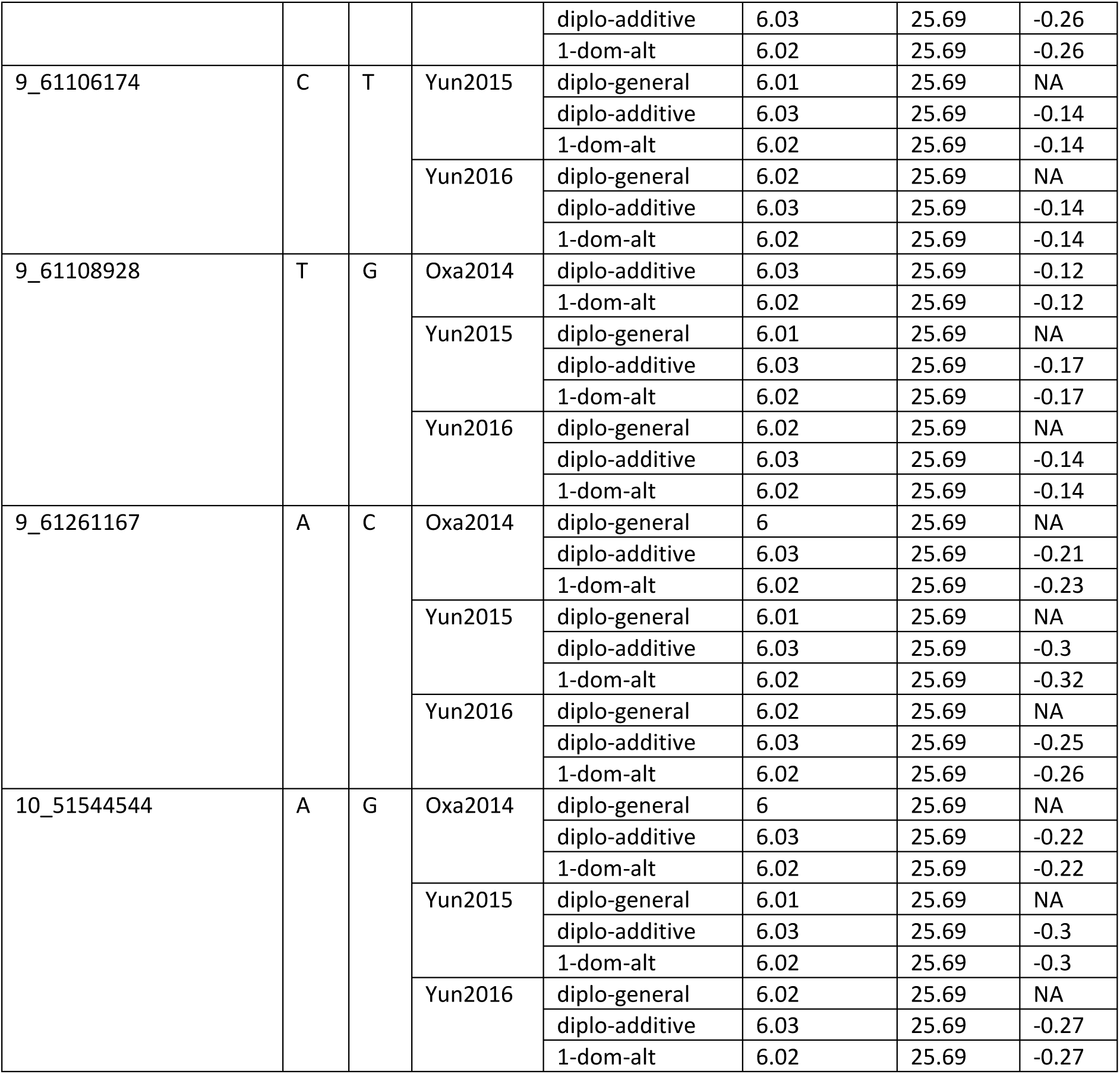
Markers tagging QTL for late blight resistance with the diplo-general, diplo-additive and 1-dominance models of GWASpoly and diploid SNP.

A few markers for late blight resistance QTL have been found in the past on chromosome III. These include a QTL tagged between the markers GP25 and CP6 (Gebhardt and Valkonen, 2001) which can be located near the locus PGSC0003DMB000000154 or approx. between 42 and 51 Mbp in the Phureja DM1-3 genome. The SNPs 3_45458723, 3_45458753 and 3_45458754 found in the diploidized dataset mapped close to this QTL. These markers are located within a WREBP-2 protein / transcription factor IIIA gene (PGSC0003DMG400009082). Marker 3_3319097 is localized within a gene encoding an iron binding oxidoreductase (PGSC0003DMG40002252) that is located near an already known QTL tagged with marker T6135 (Gebhardt and Valkonen, 2001).

Several late blight resistance genes originated from *Solanum demissum* have been already mapped to specific positions of the genome. These include *R1* in chromosome V, *R3, R6* and *R7*, all in the distal segment of chromosome XI and *R2* in chromosome IV. Also, the genes *Rber* and *Rblc* belonging to other Solanum groups have been mapped to chromosomes X and VIII, respectively. These genes show resistance to contemporary races of *Phytophthora infestans*. Additionally, promising QTL are thought to be located on chromosomes III, IV, V and VI (Gebhardt and Valkonen 2001). The region with *R1* and QTL for late blight resistance in chromosome V is flanked by the markers GP21 and GP179 approximately between 2 and 5 Mbp in the DM genome containing the SNPs 5_4260524 and 5_5572873 in the tetraploid and the diploid dataset, respectively (Table 3, 4). The SNP 10_51544544 found on chromosome X in the diploid dataset and associated with resistance in three experiments (Table 4), is mapped to a region associated with late blight resistance conferred by gene *Rber*, tagged by the marker TG63 is located approximately at 52 Mbp in the DM genome.

The SNP 6_45694949 found on chromosome VI, mapped to a physical position around 45.7 Mbp, is located near a gene and two flanking QTL that have been recently associated with quantitative late blight resistance. Álvarez et al. (2017) used association mapping to identify SNPs in genes from a set of candidate genes in *Solanum tuberosum* group Phureja associated with quantitative resistance. The gene expresses a stem 28 kDa glycoprotein and is located around 49.1 Mbp between the Ib6a and Pin6b – lb6b QTL. The favourable allele of this SNP has a significant effect on the resistance in Yunnan in 2015 and 2016, and an additive effect since the individuals homozygous for the favourable allele are more resistant than the heterozygous individuals (Figure 6).

**Figure 6.**
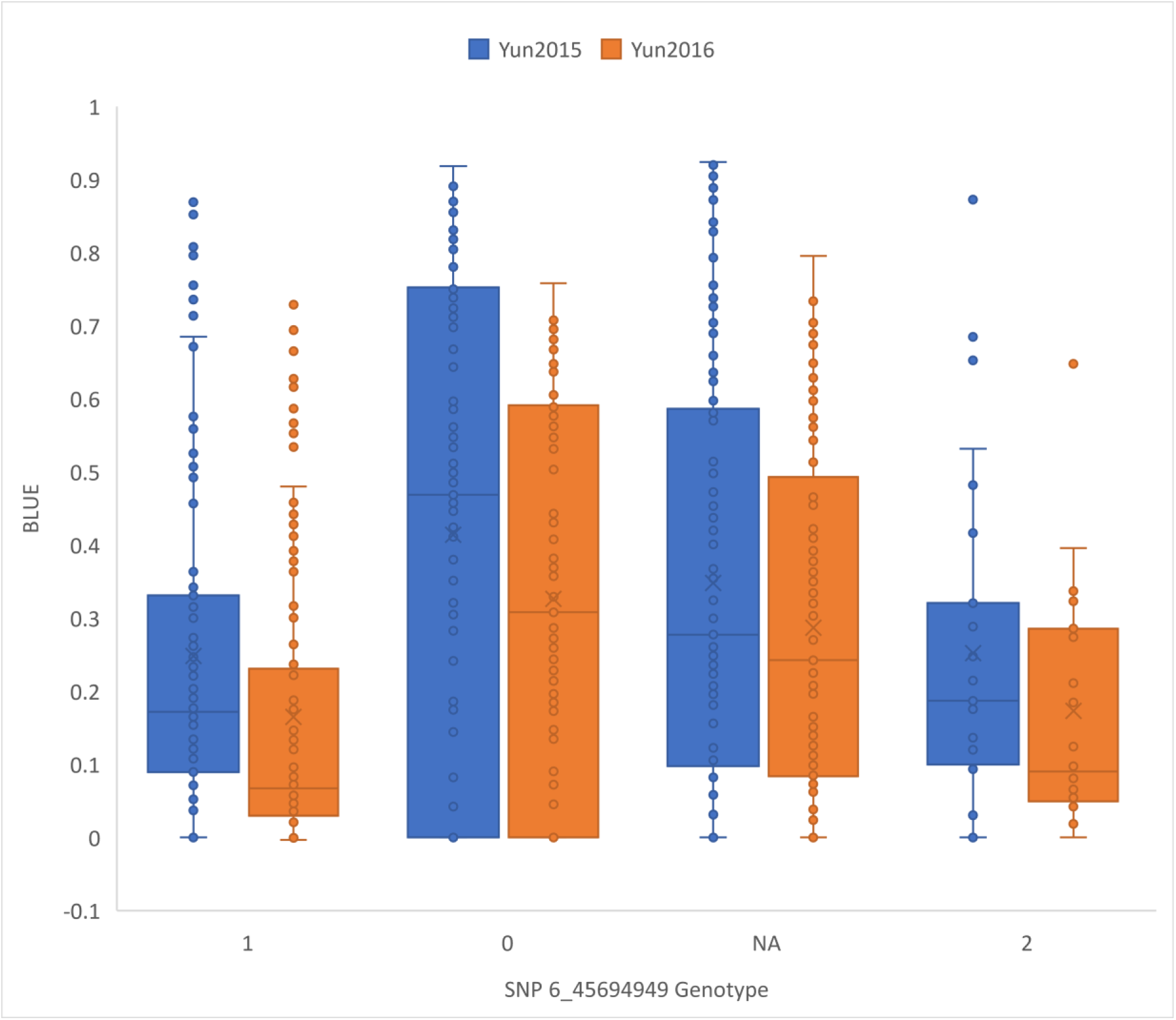
Boxplot for BLUE value distribution in the different genotype classes for the SNP 6_45694949 found in the diploidized dataset. “0” and “2” stand for homozygous for the reference and alternative alleles respectively. “1” indicates heterozygote and “NA”, missing genotype. Lower values indicate a higher resistance.

The 48x plex multiplexing of the samples during the sequencing and stringent filtering for minimum read depth in all samples yielded relatively few SNP (less than 4K), which is too few considering the level of LD decay-based estimate of 10s of thousands SNP to fully cover the genome. Therefore, the GWA was done using both tetraploid and the diplodized data. Three of the markers (9_58779951, 9_59967523, 9_60067335) map in chromosome 9 in the same chromosome region previously found associated with late blight resistance in Peru (Lindqvist-Kreuze et al., 2014; Li et al., 2010). These markers are physically separated in the DM genome by 1.3Mb, which fits the estimate for the LD decay in these potato genotypes. In the QTL dPI09c reported by Li et al. (2010) the R8 gene originating from *Solanum demissum* was recently identified (Jian Rui et al., 2018). The QTL dPI09c interval in potato DM1-3 516 R44 (Potato Genome Sequencing Consortium, 2011) begins at 60615044 bp, hence over 600Mbp away from the nearest GBS marker (9_60067335) we identified in the current research. In our tetraploid GBS marker set there are no SNP mapping in the dPI09c interval possibly because of the low sequencing depth and the complexity of the region that consists of several RXLR type resistance genes (Jiang Rui et al., 2018). In the diploid marker set, however, four markers (9_60067335, 9_61106174, 9_61108928, 9_61261167) that map in the QTL dPI09c interval were identified.

## Conclusions

## Acknowledgements

Funding from GIZ is acknowledged for the project accelerating the Development of Early-Maturing-Agile Potato for Food Security through a Trait Observation and Discovery Network. This work has also received funding from the CRP-RTB and USAID. Special thanks to CABANA project funded by Global Challenges Research Fund - part of the UK AID budget, for funding the secondment of HLK to Earlham Institute (EI) in the UK to conduct the bioinformatic analysis and to Janet Higgins at EI for the bioinformatics support.

## Contributions

Designed experiments and obtained funding: HLK, MB

Implemented field trials and collected data: MG, XL, JQ, GW, BH, ZP, QS, KN, IS

Analyzed and interpreted data: HLK, DG, BDB, PU

Bioinformatics: HLK, JDV, DG

Statistics: BDB

Wrote the manuscript: HLK, PU, MB

---

**Table S1.**
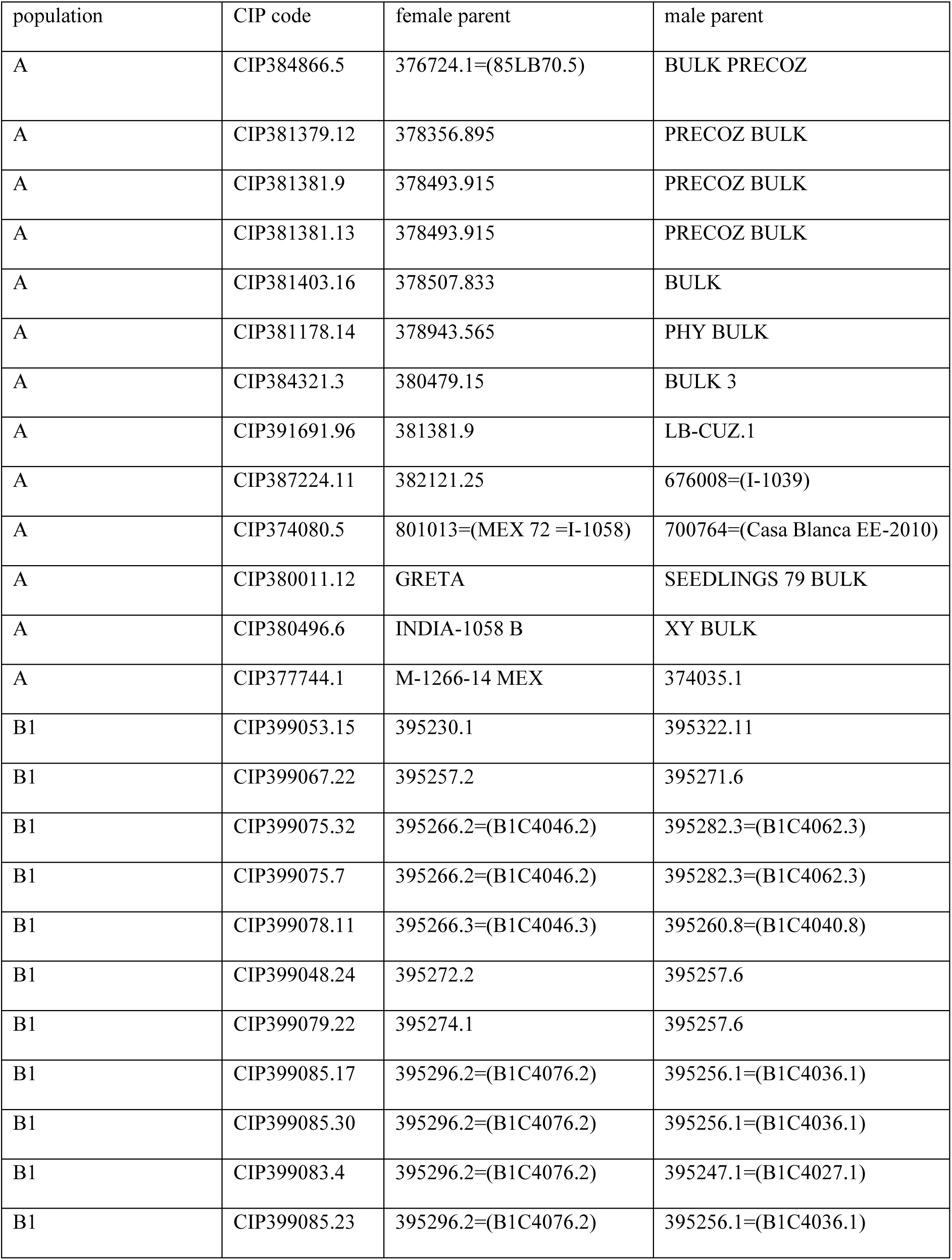
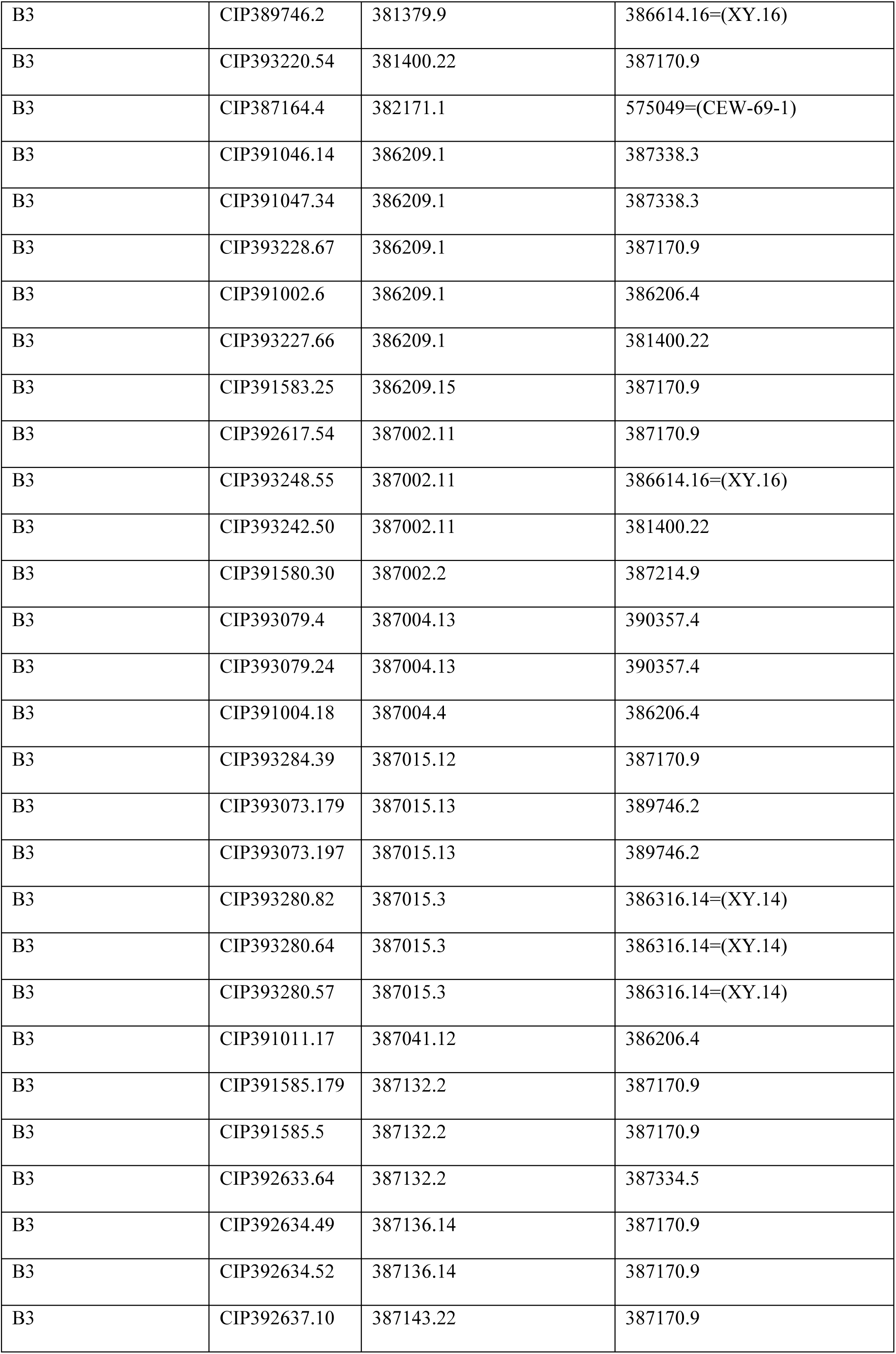
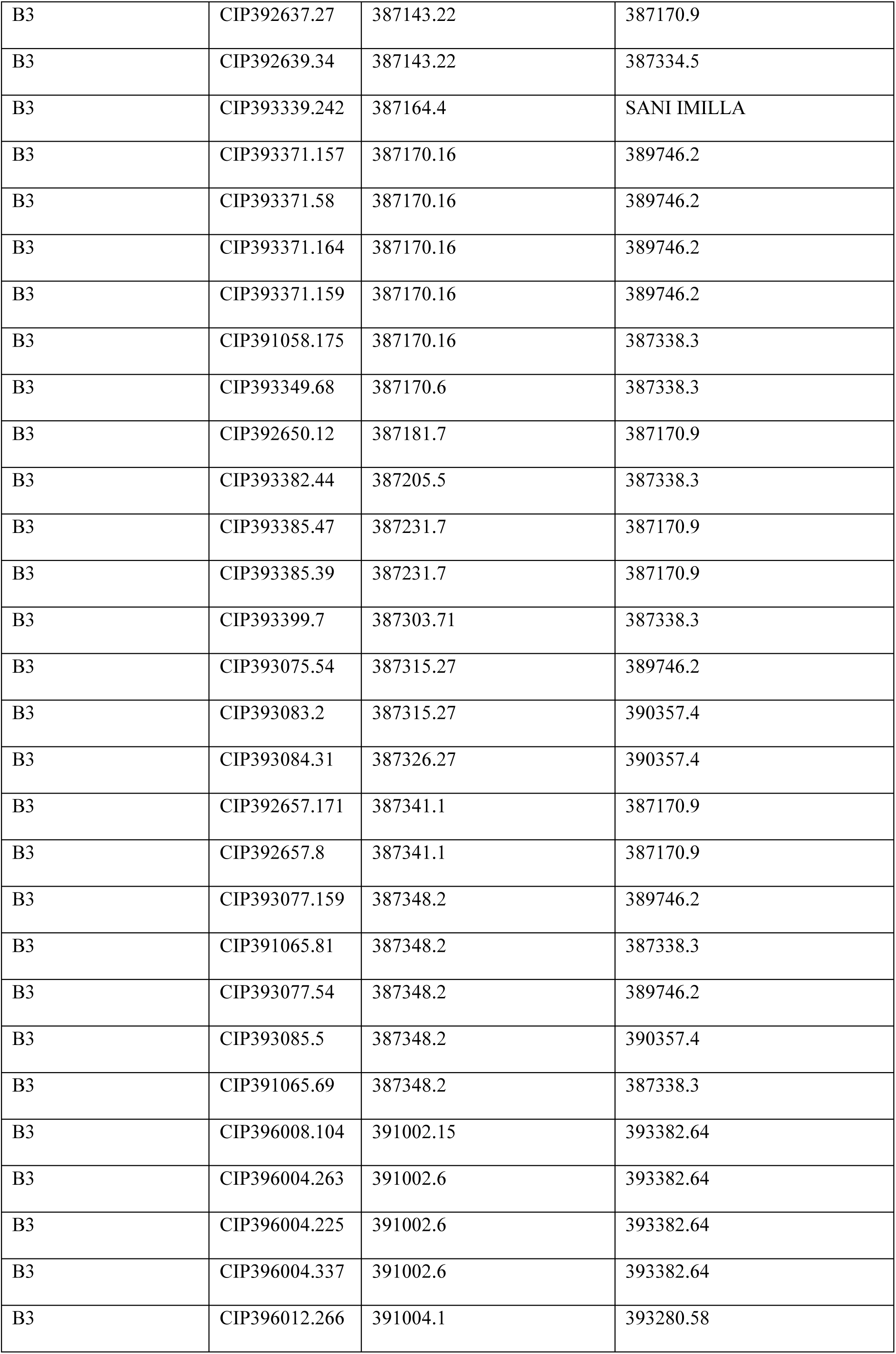
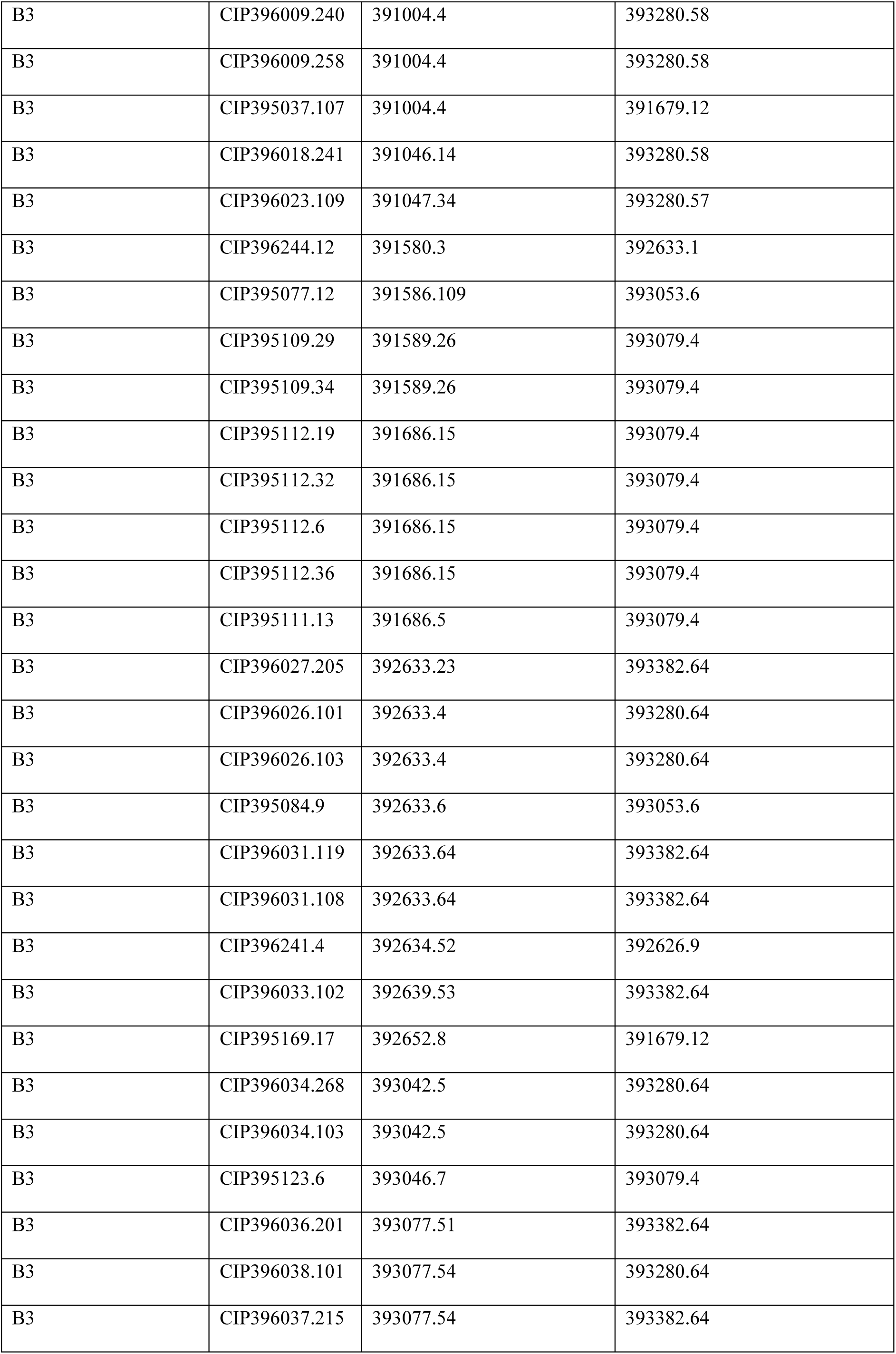
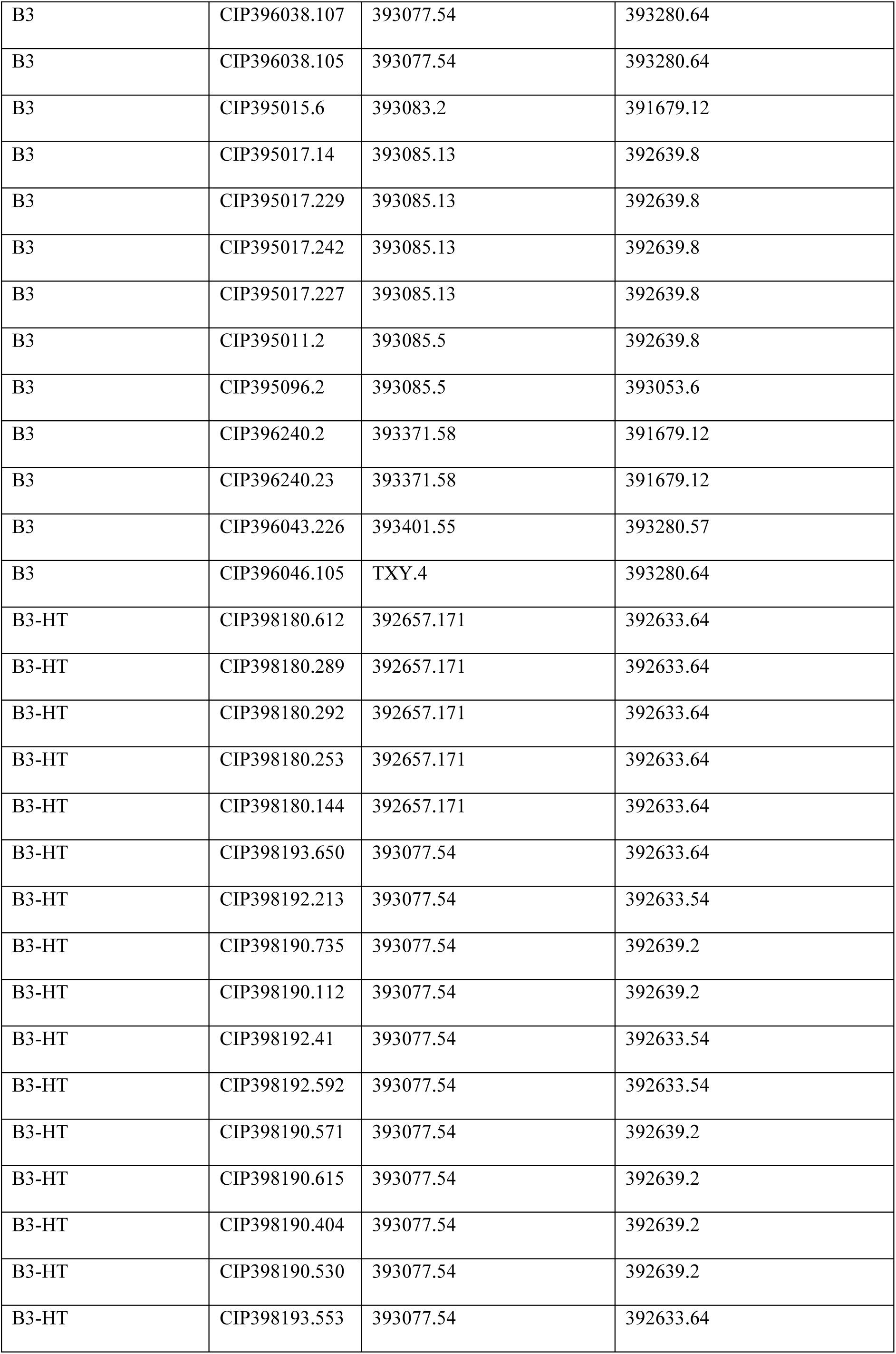
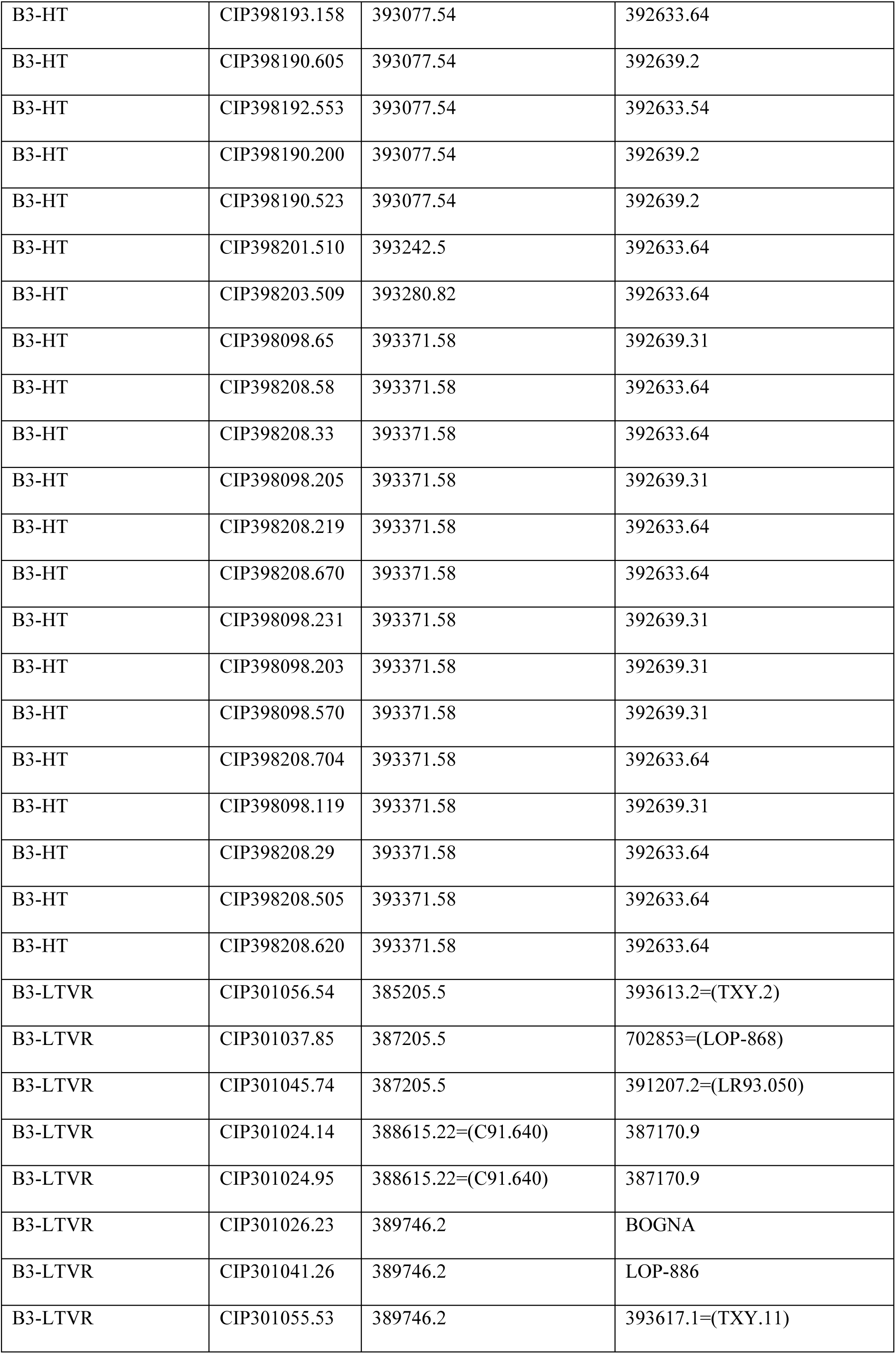
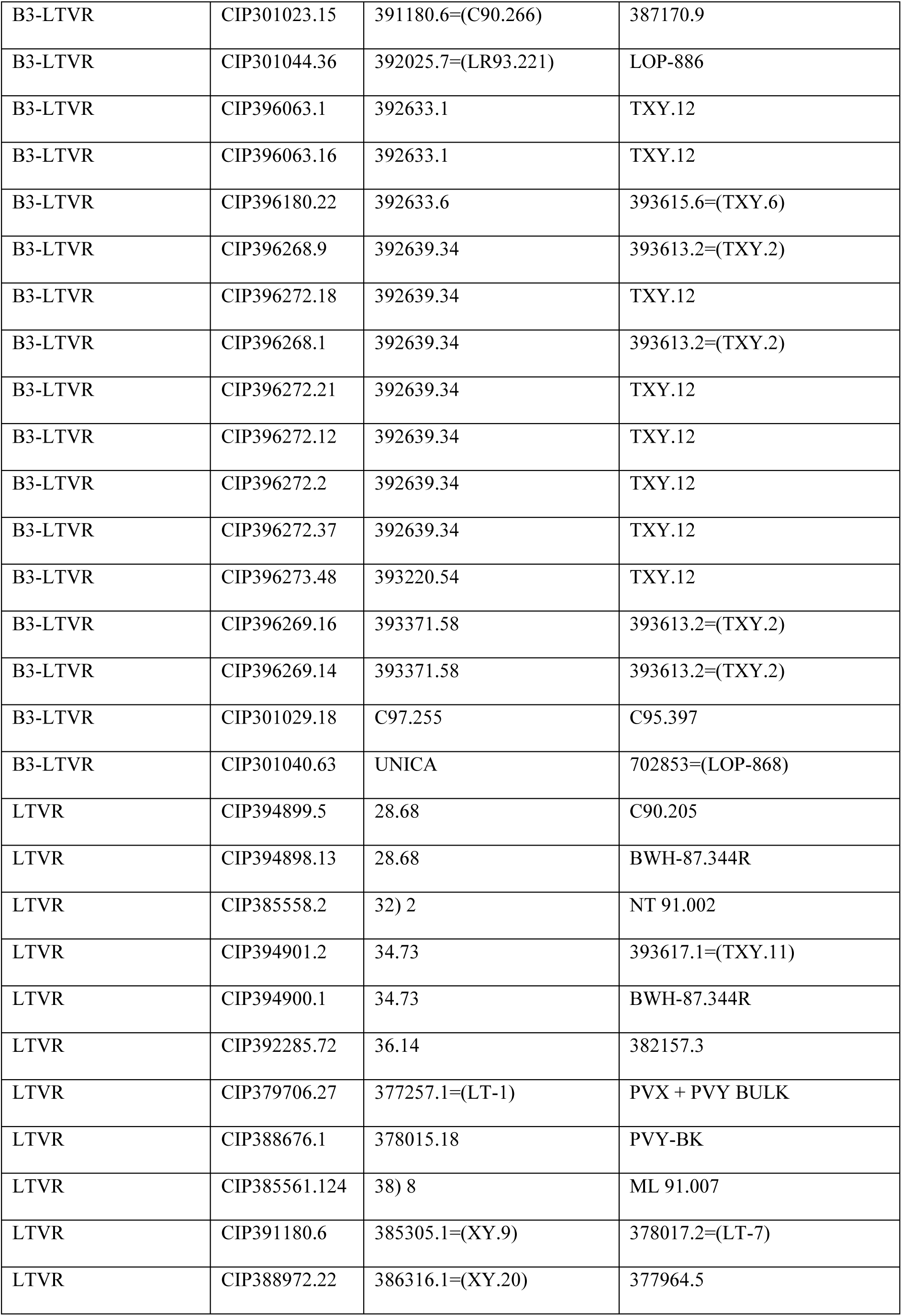
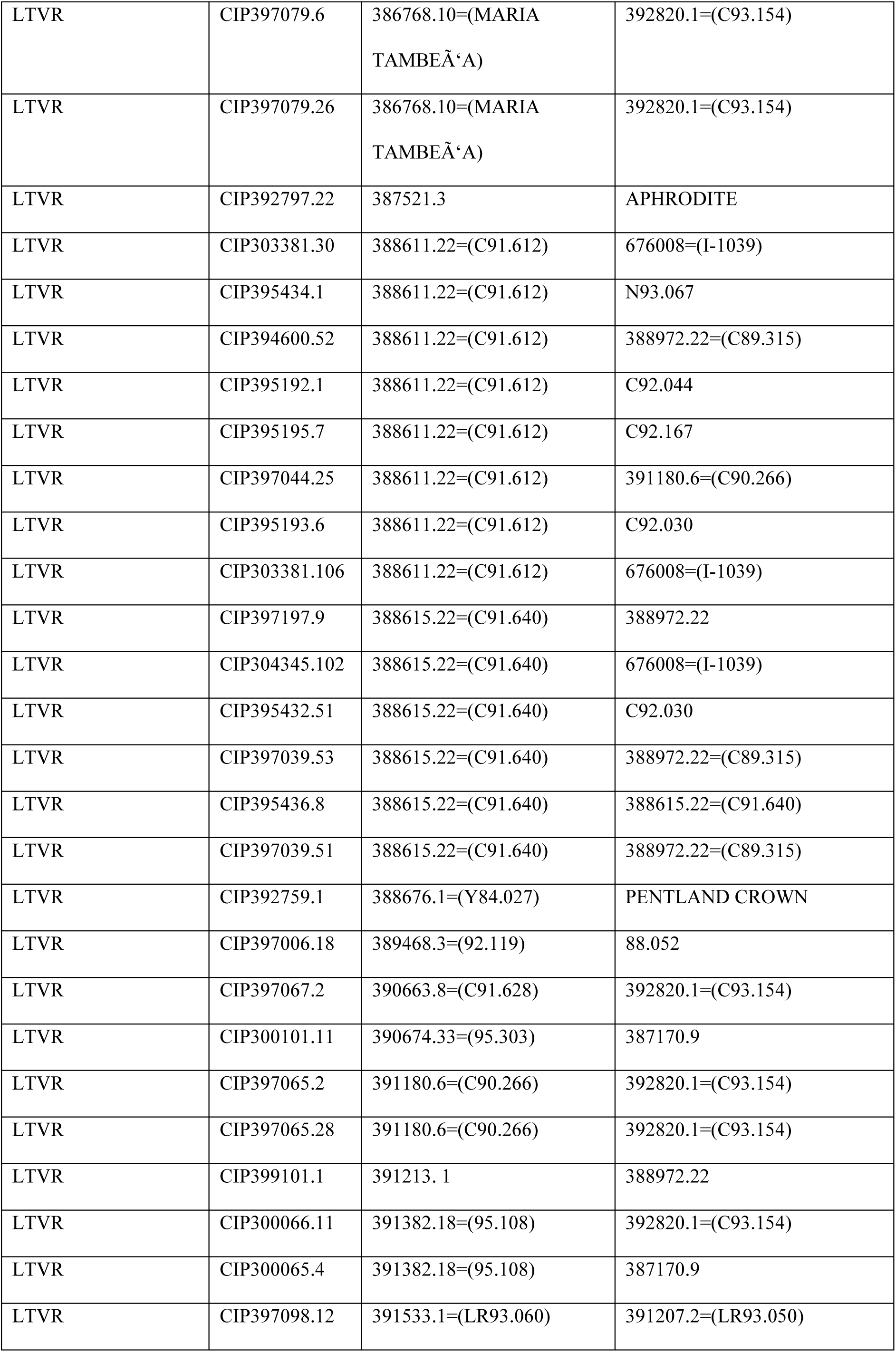
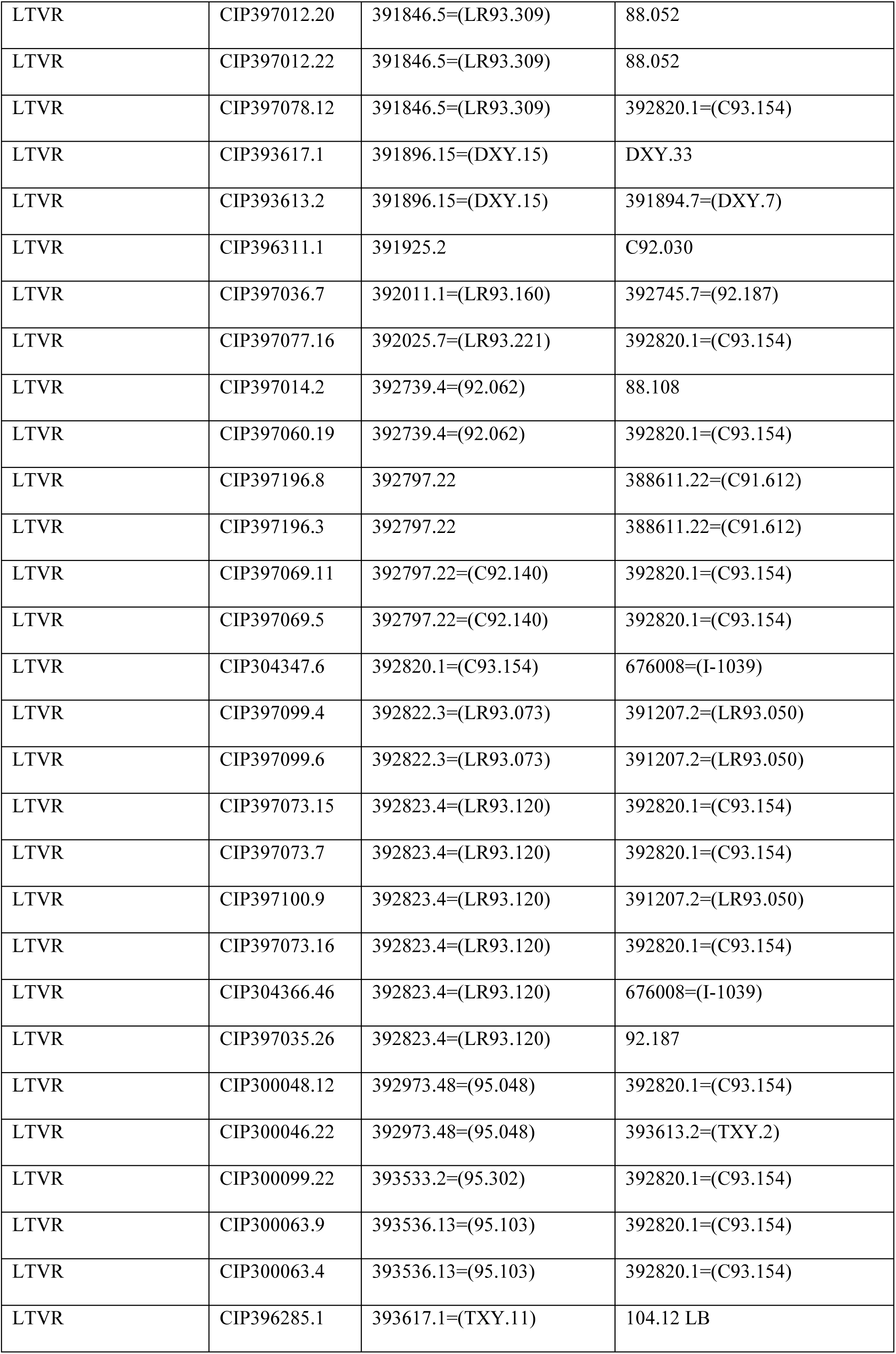
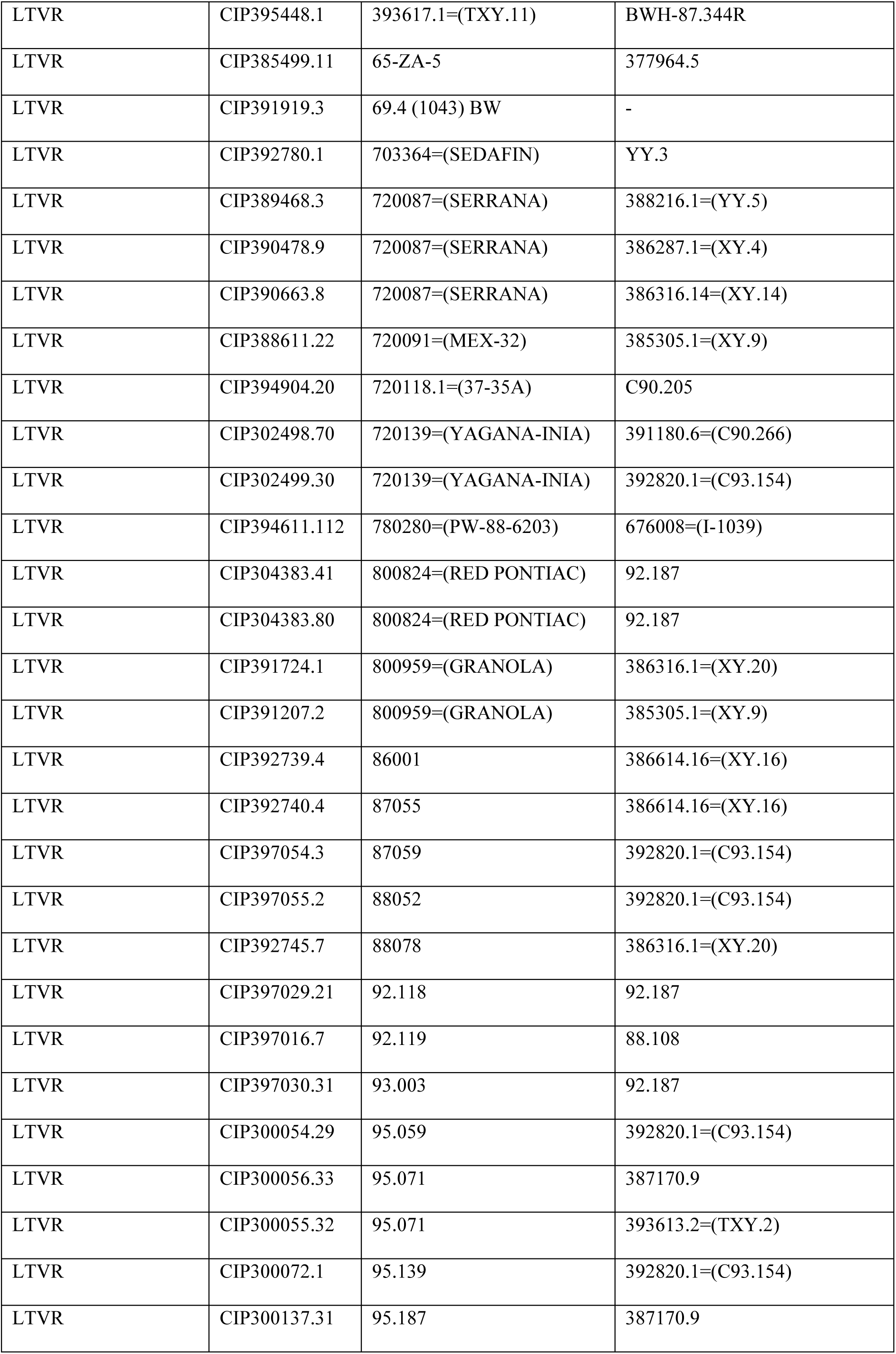
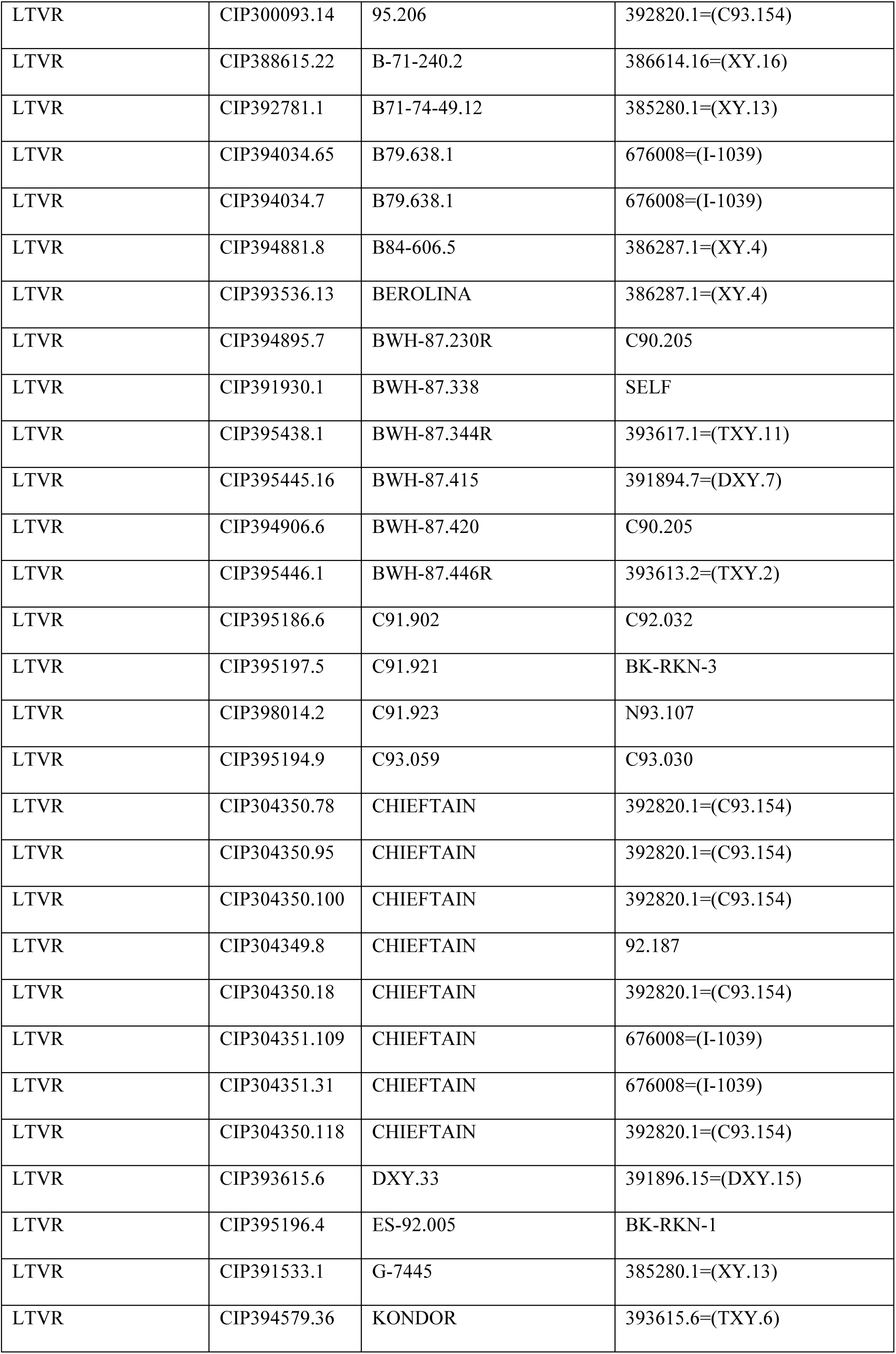
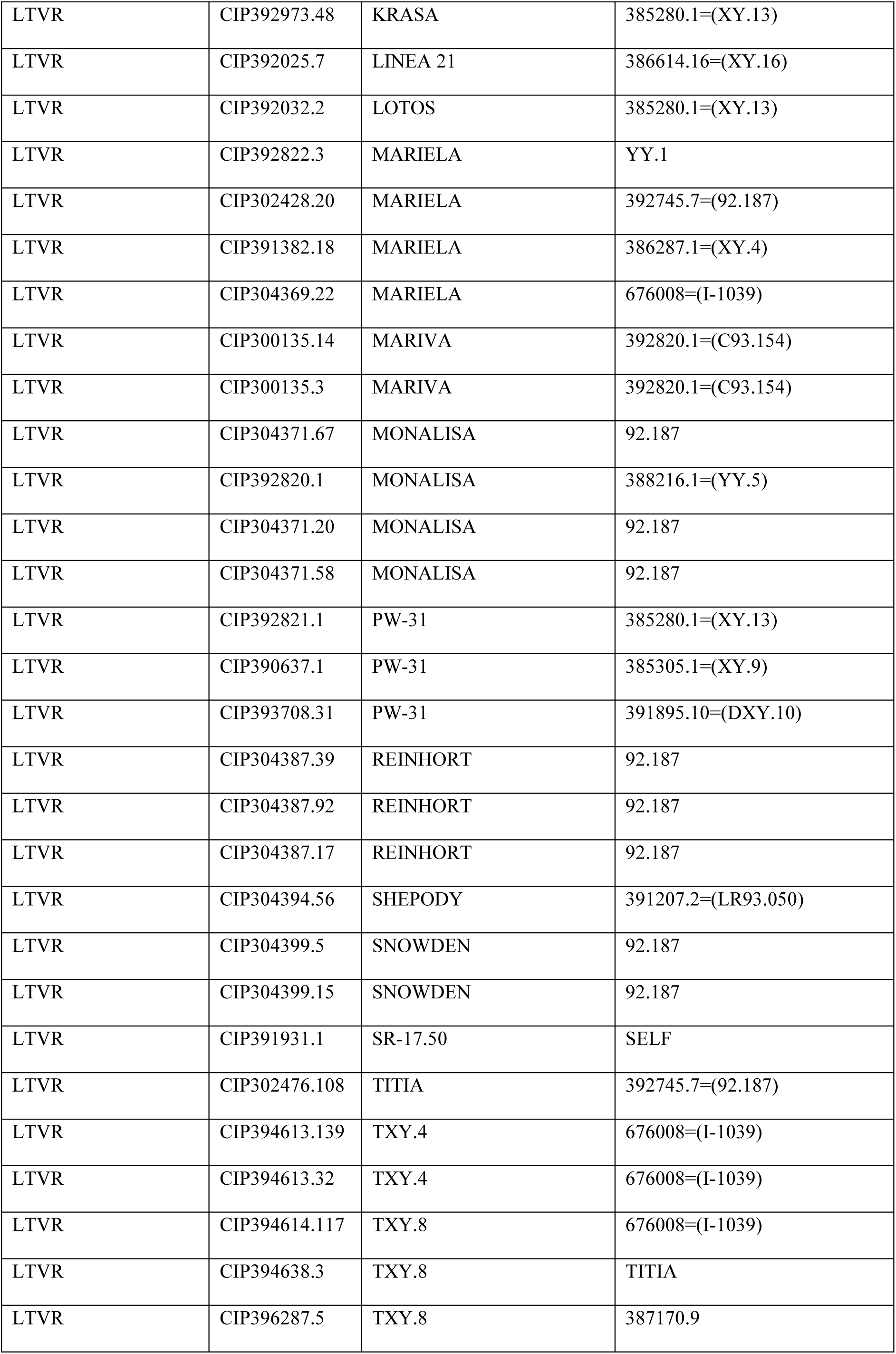
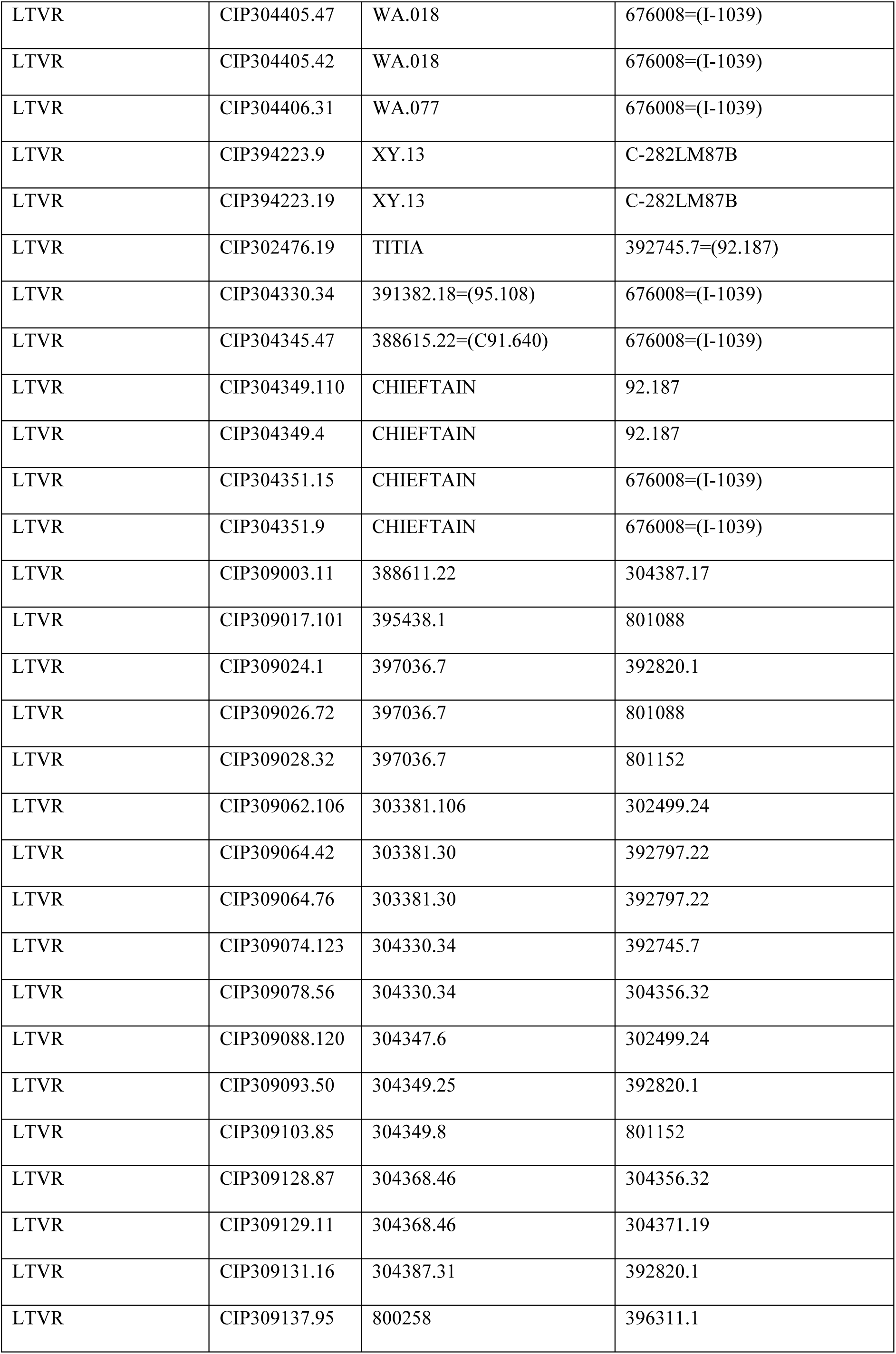
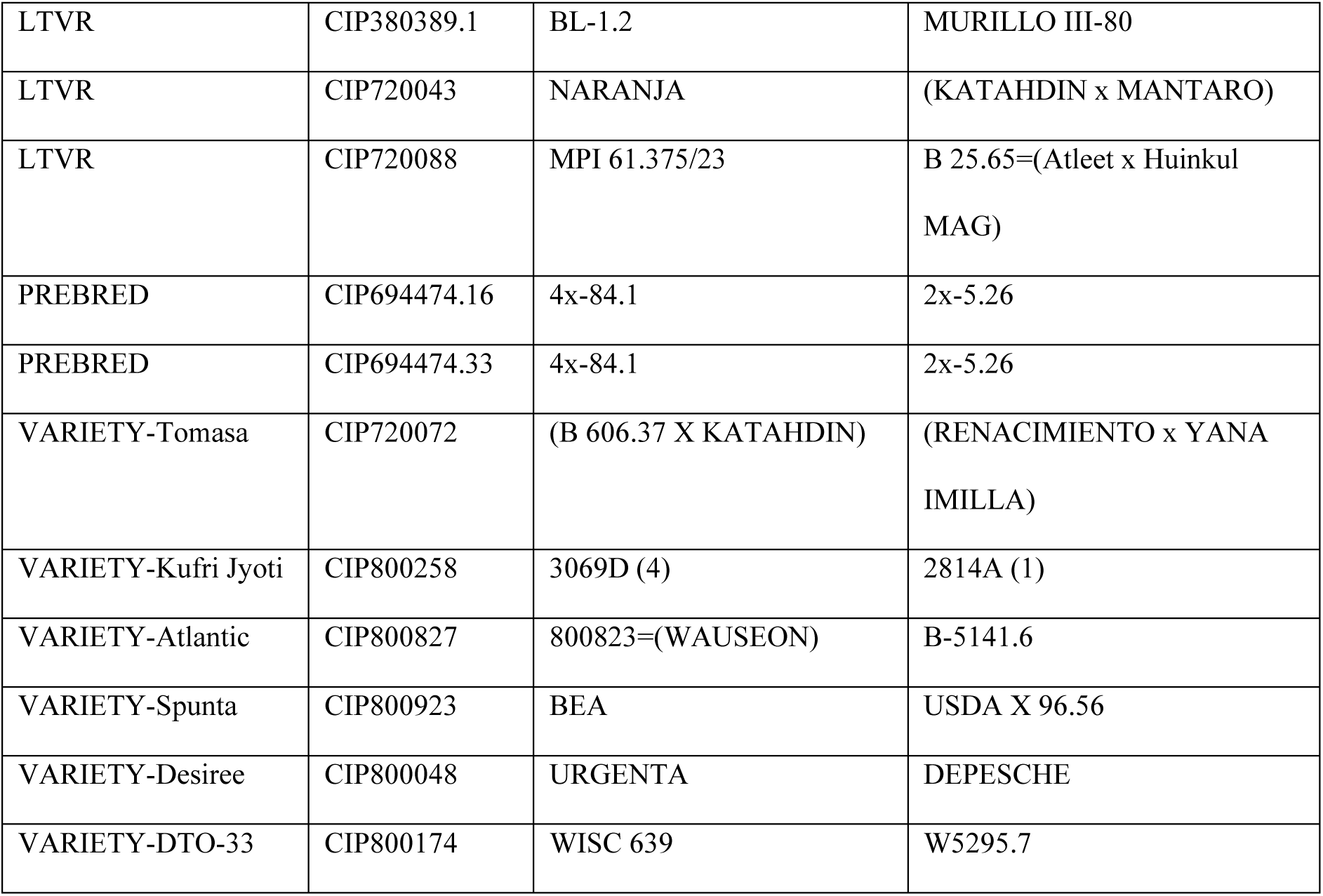
Population denominations, and parentage of the potato genotypes evaluated in this study.

